# Chilling Nights Do Not Cause Starch Over-Accumulation and Trigger a Shift in Carbon Partitioning via SPS Toward Sucrose in *Arabidopsis* - Differing from Acclimation to Permanent Cold

**DOI:** 10.64898/2026.07.31.741842

**Authors:** Christina Bonn, Oliver Giesbrecht, Angelina Jordine, Mona Becker, Sofie Jennings, Marvin van Aalst, Joost T. van Dongen, Anna Matuszyńska, Lisa Fürtauer

## Abstract

Plants acclimate to low temperatures by remodeling carbohydrate metabolism. Permanent cold at 4/5 *^◦^*C induces well-characterized cold acclimation in plants, including accumulation of starch and soluble sugars. In natural environments, however, annual plants in their vegetative phase more frequently experience chilling nights (0–6 *^◦^*C nights followed by days at least 12 *^◦^*C warmer), whose metabolic consequences remain poorly understood.

We investigated Col-0 *Arabidopsis thaliana* exposed to one or seven chilling nights and quantified central carbohydrate metabolites, starch, photosynthetic parameters, and maximal activities of enzymes linking sucrose synthesis and cleavage. We integrated these time-series data into a biologically constrained augmented neural ordinary differential equation (ANODE) model to infer diurnal reaction-rate dynamics.

Chilling nights induced strong accumulation of sucrose, glucose, and fructose, particularly after seven nights. Both measured sucrose phosphate synthase (SPS) capacity and ANODE-predicted SPS rates increased. In contrast to permanent cold acclimation, daytime starch over-accumulation was absent.

Thus, lacking starch over-accumulation together with increased SPS rates, chilling nights invoke distinct carbohydrate-partitioning patterns towards sucrose, indicating a fundamentally different acclimation strategy compared to well described permanent cold.

## 1 Introduction

Plants are naturally exposed to changing environmental conditions, such as fluctuations in light intensity, water availability, and temperature; therefore, they must fine-tune their metabolic processes in order to survive. Especially temperature fluctuations shape plant metabolism, requiring precise acclimation mechanisms to sustain physiological function (Fürtauer et al., 2019). Continuous cold (4 *^◦^*C) exposure triggers well-characterized responses typical for cold acclimation - a biological process that increases the tolerance of plants native to temperate environments to cold temperatures (C. L. Guy, 1990). This cold acclimation process involves extensive changes in gene transcription, regulatory networks and metabolism (Horton et al., 2016; Hoermiller et al., 2017; Adler et al., 2025).

On a metabolic level, carbohydrate metabolism is the key driver of cellular adaptation, as it rapidly provides energy (e.g. adenosine triphosphate (ATP)) to support physiological stress responses and gene expression (Rosa et al., 2009). Furthermore, its metabolic intermediates function as essential signaling molecules and biosynthetic building blocks that directly guide cells to adapt to and survive changing environmental conditions (Thalmann & Santelia, 2017; Saddhe et al., 2020).

A clear distinction between the immediate stress response, when a plant is exposed to the colder temperatures for the first time (cold shock), and the acclimated state, which a plant enters after a prolonged cold period over several days (cold acclimation), was found (Huner et al., 1998; C. Guy, 1999; Nägele & Heyer, 2013; Hoermiller et al., 2017; Zuther et al., 2018; Dong & Beckles, 2019; Fürtauer et al., 2019; Herrmann et al., 2019; Kitashova et al., 2023).

The immediate stress response, within the first 24 h after the stress is introduced, heavily impacts sucrose cycling, i.e. cyclic biosynthesis and degradation of sucrose (Weiszmann et al., 2018; Fürtauer et al., 2019). Sucrose cycling is ought to stabilize metabolism during stress, even if itself seems to be energetically wasteful (futile cycle), but it allows precise control over carbon partitioning (Ruan, 2014). For immediate, permanent cold stress, this includes accumulation of soluble sugars such as sucrose, glucose and fructose as well as increased enzyme activities of glucokinase (phosphorylation of glucose to glucose-6-phosphate), neutral invertase (hydrolysis of sucrose to glucose and fructose) while fructokinase (phosphorylation of fructose to fructose-6-phosphate) remained stable (Herrmann et al., 2019; Kitashova et al., 2023). It was shown that sucrose phosphate synthase (sucrose synthesis from fructose-6-phosphate and uridindiphosphat-glucose) over expression lines lead to an increase in photosynthetic activity and therefore increased freezing tolerance (Å. Strand et al., 2003; Herrmann et al., 2019). At the same time in studies to permanent cold acclimation no significant increase of sucrose phosphate synthase maximum activity has been reported (Kitashova et al., 2021, 2023). Furthermore starch degradation via amylase is up-regulated resulting in lower starch concentrations and consequently a supply of glucose to the energy metabolism (Thalmann & Santelia, 2017; Kitashova et al., 2023). Such immediate stress responses provide the energy and resources required for the synthesis of new metabolites involved in plant protection, repair, and acclimation (Ribeiro et al., 2022).

During the acclimated state, on the other hand, starch accumulates as part of an adaptive strategy to maintain carbohydrate homeostasis (Thalmann & Santelia, 2017; Ribeiro et al., 2022; Kitashova et al., 2023) and under hypothermia granules serve as photoprotectors (Popov, 2026). Specifically, in several plant species including *Arabidopsis*, starch and soluble sugars like sucrose, glucose and fructose are reported to highly accumulate during permanent cold to serve as energy reserves (Scarth & Levitt, 1937; Stitt & Hurry, 2002; Klotke et al., 2004; C. Guy et al., 2007; Nagler et al., 2015; Garcia-Molina et al., 2020; Kitashova et al., 2023; Popov, 2026) as well as osmoprotectants, membrane stabilizers and desiccation protectors (Zuther et al., 2018). This is accompanied by a finely tuned modulation of enzyme activities, whereby certain enzymes (such as fructokinase) are up-regulated, while others (e.g. invertases) return to a more stable level after an immediate response in order to control the balance between soluble sugars and starch accumulation (Pommerrenig et al., 2018; Kitashova et al., 2021, 2023). Both immediate and acclimated changes in the carbohydrate metabolism contribute to an increased freezing tolerance of the plant (Scarth & Levitt, 1937; A. Strand et al., 1999; Nägele & Heyer, 2013; Fürtauer & Nägele, 2016; Weiszmann et al., 2017; Fürtauer et al., 2019; Kitashova et al., 2021; Seydel et al., 2022).

In natural habitats, however, annual plants are rarely exposed to constant cold temperatures. Instead, they experience temperature fluctuations, which can be particularly high between day and night (up to 20 *^◦^*C (Wetterdienst, 2026)). These diurnal temperature differences have been shown to get more extreme in temperate and polar regions (Liu et al., 2024). “Chilling nights”, where the temperature decrease towards low temperatures above the freezing point (between 0 *^◦^*C and 6 *^◦^*C) in the night, followed by moderate day temperatures (at least 12 *^◦^*C higher than the minimal temperature), occur naturally in spring and autumn (Unterberger et al., 2018). Due to climate change, the difference between minimum night temperatures and maximum day temperatures as well as the occurrences of chilling nights increased in the past years (G. Wang & Dillon, 2014; Liu et al., 2024) and are exemplarily shown for Central Europe Germany and Poland, in Suppl. Fig. S1. Those chilling nights affect annual plants of temperate regions, including *Arabidopsis thaliana* precisely when the plant reaches its vegetative stage. Despite their ecological relevance, the metabolic consequences of chilling nights have barely been examined. To fill this gap, we investigated how chilling nights influence the central carbohydrate metabolism in *A. thaliana*.

We investigated whether the perception and response of *A. thaliana* Columbia-0 (Col-0) to intermittent cold differs from its response to prolonged low temperature exposure using a time-series analysis. By determining carbohydrate concentrations of six carbohydrates (glucose-6-phosphate (G6P), fructose-6-phosphate (F6P), sucrose (suc), glucose (glc), fructose (frc), and starch), and the maximum activities (*v* _max_) of key enzymes involved in sucrose metabolism (sucrose phosphate synthase (SPS), acidic and neutral invertase (aINV and nINV), glucokinase (GLCK) and fructokinase (FRCK) (Fig. 1A) and integrated these data into an augmented neural ordinary differential equation (ANODE) model (Gholami et al., 2019). This approach enabled us to infer continuous metabolic flux trajectories directly from the time-series data, without requiring a fully specified kinetic model. It is particularly suited to carbohydrate metabolism under chilling stress, where the underlying regulatory state remains incompletely characterized and is only partially captured by the measured metabolites. By combining experimental metabolite and enzyme activity measurements with ANODE-based modeling, we reconstructed dynamic metabolic fluxes across the diurnal cycle and identified a distinct acclimation strategy under chilling-night conditions.

**Figure 1:**
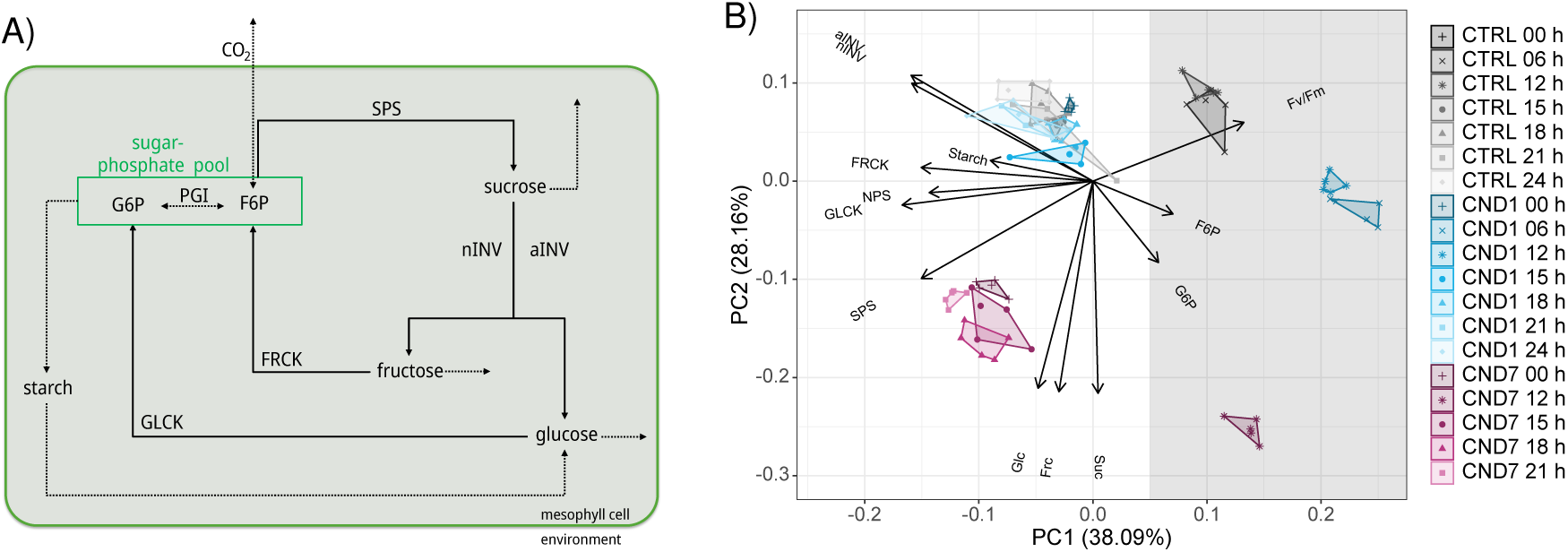
Model of metabolism and PCA of all measured data. **A)** Simplified carbohydrate metabolism representative for the used model including CO_2_ fixation/release and formation/degradation of sugar phosphates (via carbon exchange), suc synthesis (via SPS), starch accumulation, and suc degradation by INV followed by phosphorylation of glc and frc by GLCK and FRCK, respectively, leading to F6P and G6P which are interconverted reversibly by PGI, with export of suc to sinks (Nägele et al., 2010, 2012; Fürtauer & Nägele, 2020; Giesbrecht et al., 2025). Solid arrows: modeled rates with measured *v* _max_ values, dashed lines: modeled reaction rates without *v* _max_ input. **B)** Principal component analysis for all gathered NPS, F_v_/F_m_, metabolite and Arrhenius corrected enzyme data by time point and condition. CTRL data are depicted in shades of gray, CND1 data in shades of blue and CND7 data in shades of magenta. The gray background on the right side indicates the separation of the night from the day on the left. Abbreviations: CND1:= first chilling night 4 *^◦^*C/22 *^◦^*C night/day; CND7:= seventh chilling night 4 *^◦^*C/22 *^◦^*C night/day; CTRL:= control 18 *^◦^*C/22 *^◦^*C night/day; F6P:= fructose-6-phosphate; frc:= fructose; FRCK:= fructokinase; Fv/Fm:= maximum potential quantum efficiency of Photosystem II (PSII); G6P:= glucose-6-phosphate; glc:= glucose; GLCK:= glucokinase; n/aINV:= neutral/acidic invertase; NPS:= net photosynthesis; PGI:= phosphogluco isomerase; SPS:= sucrose phosphate synthase; suc:= sucrose.

Dissecting intermittent temperature stress is essential for a realistic understanding of natural plant cold tolerance mechanisms. Our results showed that during chilling nights (i) soluble sugar levels and (ii) SPS rates (quantified *v* _max_ and modeled *v* _pred_) increase strongly, (iii) starch does not accumulate during the day under chilling night conditions as it does under permanent cold and (iv) FRCK rates are significantly reduced. Consequently, permanent cold and chilling nights trigger fundamentally different acclimation strategies with a shift towards sucrose metabolism and lack of starch accumulation in chilling nights.

## 2 Materials and Methods

### 2.1 Plant Material and Growth Conditions

*A. thaliana* Col-0 seeds (NASC ID: N22660 (Nordborg collection); N22625)) were stratified dark in water for three days at 4 *^◦^*C. Subsequently, seeds were sown onto expanding biodegradable turf plant pots (Jiffy pellets (7) 44 mm (turf coco)) and cultivated for 37 days in a GEN1000 Conviron (Conviron, Canada). Plants were grown in a 12 h night / 12 h day cycle, with 18 *^◦^*C nights and 22 *^◦^*C days. Humidity was set to 60% and light intensity to 100 µmol m*^−^*^2^ s*^−^*^1^, including a 30 min dusk dawn transition phase.

After 37 days, a chilling 4 *^◦^*C night was induced, while the temperature during the day was kept at 22 *^◦^*C to recreate realistic conditions plants face in spring and autumn in temperate regions. The transition from 22 *^◦^*C to 4 *^◦^*C took 1 h 20 min and 55 min vice versa. Control plants (CTRL) and chilling night plants (CND1) were harvested during the first 24 h after the first cold night was induced: at the beginning of the night (00 h), the middle of the night (06 h), the end of the night (12 h) and every three hours during the following day at the 15 h, 18 h, 21 h, and 24 h time points.

In addition, samples were harvested on the seventh day (CND7) when plants had already been exposed to six 4 *^◦^*C nights each followed by a 22 *^◦^*C day at the 00 h, 12 h, 15 h, 18 h and 21 h time points. Harvested plants were immediately frozen in liquid nitrogen, ground to fine powder and freeze dried.

For the determination of the maximum invertase activities and hexosephosphate concentrations, a second batch of plants was not lyophilized – the batches were compared through the measurement of metabolite concentrations at selected time points and were determined not to be significantly different. The ratio between fresh mass (FM) and dry mass (DM) was determined to be 11.3 *±* 0.4. Net photosynthesis and F_v_/F_m_ measurements were performed on living plants.

### 2.2 Net Photosynthesis and Capacity of Photosystem II

Net photosynthesis and F_v_/F_m_ measurements were performed with a GFS-3000 (WALZ®, Effeltrich, Germany), infrared gas analyzer. Day samples were 10 min dark acclimated before the F_v_/F_m_ determination via a pulse-amplitude-modulation (PAM) measurement and subsequently net photosynthesis was determined after signal stabilization (approximately 10 minutes) via CO_2_ and H_2_O measurements. For the night samples, first CO_2_ and H_2_O measurements were performed followed by F_v_/F_m_.

### 2.3 Metabolite Analysis

#### 2.3.1 Sucrose, Glucose and Fructose Quantification

The soluble sugars were extracted from ∼2.5 mg dried leaf material by double ethanolic extraction (80%) at 80 *^◦^*C for 30 min (Suppl. A.2.1). The extract was dried and sugars were resolved in ddH_2_O and the pellet further used for starch quantification. For sucrose (suc) quantification, monosaccharides were removed by incubation with 30% KOH solution at 95 *^◦^*C for 10 min. Subsequently, the amount of suc was determined via anthrone assay (adapted from Dreywood (1946), Suppl. A.2.1). For the quantification of glucose (glc) and fructose (frc) a coupled enzyme assay was performed. In an reaction cascade with phosphoglucoisomerase (PGI), hexokinase (HXK) and glucose-6-phosphate dehydrogenase (G6PDH) reduced nicotinamide adenine dinucleotide phosphate (NADPH) is formed and measured at 340 nm ((Schmidt, 1961), Suppl. A.2.1). For all sugars, standards were determined in parallel and absolute amounts were determined in the samples.

#### 2.3.2 Starch Quantification

The remaining pellet of the ethanolic extraction (see 2.3.1 and Suppl. A.2.1) was used for starch quantification as described by Chow and Landhausser (2004) with alterations described in Suppl. A.2.1.

#### 2.3.3 G6P and F6P Quantification

To quantify sugar phosphates, glucose-6-phosphate (G6P) and fructose-6-phosphate (F6P) were extracted from ground plant material (fresh mass, FM) using a TCA-extraction protocol (see Suppl. A.2.1), which was adapted from Jelitto et al. (1992) and Hajirezaei et al. (2003). Then, a cycling assay was performed as described by Gibon et al. (2002), based on the findings of Nisselbaum and Green (1969), with slight alterations described in Suppl. A.2.1 to quantify G6P and F6P.

### 2.4 Quantification of Enzyme Activities

#### 2.4.1 SPS Maximum Activity

The assay to determine the maximum SPS activity was based on the assay described by J. L. A. Huber et al. (1989) with alterations described in Suppl. A.2.2. After incubation suc formation is stopped by addition and incubation with KOH for the removal of contained monosaccharides, to enable suc quantification via the Anthrone assay as described above (see 2.3.1).

#### 2.4.2 INVs Maximum Activities

With this assay, *v* _max_ of the soluble acidic invertase (aINV) and the neutral invertase (nINV) were determined as previously described by Sung et al. (1989) with alterations (Suppl. A.2.2). Enzymes were extracted using a detergent-based extraction buffer. Subsequently, for aINV and nINV, activity samples were incubated with acidic or neutral reaction buffer respectively. After neutralization, the reaction was heat inactivated. The amount of glc was then determined as described for the starch quantification assay (Suppl. A.2.1).

#### 2.4.3 HXKs Maximum Activities

The maximum hexokinases activities were determined with the assay described by Wiese et al. (1999) with slight alterations (Suppl. A.2.2). Maximum activities *v* _max_ were then determined based on the resulting slopes of the kinetic measurements and the law of Lambert-Beer (Wypych, 2015).

### 2.5 Arrhenius Correction

All enzyme activities were measured at the optimal temperature for each enzyme in the laboratory. To consider the actual temperatures the plants faced at each of the corresponding time points, the Arrhenius-equation was used (Arrhenius, 1889a, 1889b; Laidler, 1984). The 00 h time point was considered as 22 *^◦^*C since plants were exposed to this temperature for 12 h before the harvest. Solely the 06 h and 12 h time points were corrected to the corresponding nighttime temperatures: 18 *^◦^*C in control and 4 *^◦^*C in chilling night day 1/7. Used activation energy of each enzyme (Kitashova et al., 2023) as well as all the pre-Arrhenius-corrected *v* _max_ values can be found in the supplementary data (Table S1).

### 2.6 Data Analysis

All statistical methods as well as the principal component analysis (PCA) and correlation matrices were conducted with R 4.2.3 in R-Studio (R Core Team, 2023). To statistically analyze the measured data, t-tests, one-way ANOVAs (analysis of variance), and Tukey’s HSD (honestly significant difference) test were conducted. The results obtained by t-test are depicted by asterisk while the output of a Tukey’s test is depicted with a compact letter display (CLD) (Abdi & Williams, 2010).

### 2.7 Mathematical Model and Numerical Simulations

A neural ordinary differential equation (NODE) framework (Chen et al., 2019) was implemented using the mxlpy Python package (Kidger, 2021; van Aalst et al., 2025), with an augmented latent state following the ANODE formulation, to account for unobserved biochemical factors (Dupont et al., 2019) (Suppl. A.2.3). The dynamics of the observed six metabolites were constrained by a fixed stoichiometric representation of the core carbon metabolic network, while the underlying reaction fluxes and latent-state evolution were learned from the data using fully connected neural networks. Additional experimentally measured enzyme activities were incorporated as time-dependent constraints on the corresponding fluxes, enabling the model to combine prior biological knowledge with data-driven flexibility.

Model parameters were optimized by minimizing the discrepancy between predicted and observed metabolite trajectories using a curriculum-based training procedure designed to improve the stability of long-horizon ODE optimization. To account for sensitivity to random initialization, an ensemble of 100 independently trained models was generated for each experimental condition (control, chilling night day 1, and chilling night day 7), and reported metabolite and flux predictions (*v* _pred_) correspond to the ensemble mean *±* SEM (standard error of the mean) (see Suppl. A.2.7).

## 3 Results

### 3.1 Night-Time Separation on Day 1 and Clear Differentiation after 7 Repeated Chilling Nights

To answer the question of how plants adapt and acclimate to chilling night conditions - nocturnal temperatures of 0-6 *^◦^*C followed by daytime temperatures *≥*12 *^◦^*C higher - we examined the ecotype Col-0 of *Arabidopsis thaliana* in a time series, at the first and seventh day of seven consecutive chilling night events (4 *^◦^*C night / 22 *^◦^*C day). We determined main carbohydrate concentrations of starch, glucose-6-phosphate (G6P), fructose-6-phosphate (F6P), sucrose (suc), glucose (glc), fructose (frc) and the maximum activity (*v* _max_) of their interconnecting enzymes of sucrose phosphate synthase (SPS), acidic and neutral invertase (aINV and nINV), glucokinase (GLCK), and fructokinase (FRCK) as well as photosynthetic activities (Fig. 1A and 1B).

In the principal component analysis (PCA) of the time-series data, time points in the middle and at the end of the night (*×* 06 h and *∗* 12 h) already separated clearly under control conditions (CTRL, shades of gray) from time points of the light phase and the beginning of the dark phase along principal component one (Fig. 1B, gray vs. white background).

The first chilling night (chilling night day 1, shades of blue) led to a strong initial stress reaction, separating the night time points even further from the day time points, which remained in the range of the control day time points. Exposure to seven consecutive chilling nights (chilling night day 7, shades of red) led to a decrease in the distance between the night and day time points. At the same time, chilling night day 7 samples were generally clearly separated from both control and chilling night day 1 (Fig. 1B magenta vs. gray and blue).

The loadings in the PCA showed that the main differentiation (∼38%) between the night and day time points was mainly driven by the enzymes GLCK and FRCK, net photosynthesis and starch. The separation of the acclimated day 7 plants compared to the day 1 plants on the second principal component (∼28%) was mainly due to the soluble sugars suc, glc and frc and to some extent SPS.

In summary, the PCA revealed a strong day–night separation, an enhanced nighttime stress response on day 1, and a clear shift toward a distinct acclimated state after seven chilling nights (Fig. 1B).

### 3.2 Constrained ANODE Modeling Provides Ensemble-Based Estimates of Reaction-Rate Dynamics

A standard ANODE learns the entire system dynamics in an abstract, “black-box” space, so the quantities it recovers don’t necessarily correspond to real, physiologically meaningful reaction rates. To avoid this interpretability problem the stoichiometry of the metabolic network was fixed beforehand and the neural network only learned the fluxes through it, so that estimated fluxes do correspond to genuine reaction rates. The functional dependence of each flux on the underlying metabolite concentrations remains learned rather than mechanistic, however, so while the recovered flux magnitudes are biologically interpretable, the kinetics and regulation that produce them are not.

To keep the fitted fluxes within a biologically plausible range despite this, we additionally bounded reaction rates by their experimentally measured *v* _max_. Supplementary, an ensemble modeling strategy was employed, refitting the model independently multiple times and retaining only those fits that met a predefined accuracy threshold (see methods 2.7). Each retained fit in the ensemble represents an equally valid explanation of the observed sugar and starch dynamics under the assumed network topology. The spread of flux values across the ensemble at each time point reflects the uncertainty in the underlying flux combinations, conditional on the fixed stoichiometry and model architecture. Reporting the mean and standard deviation of the ensemble fluxes provides a more robust estimate than a single trajectory would.

### 3.3 Both SPS Maximum Enzyme Activity and Predicted Reaction Rates are Elevated in Chilling Night Day 7

To investigate the carbon balance we looked into the sucrose cycling starting with sucrose phosphate synthase (SPS). Substrates sugar phosphates (F6P, uridine diphosphate-glc (UDP-glc)) yield UDP and sucrose 6-phosphate via SPS, which is followed by sucrose phosphate phosphatase (SPP) to gain sucrose, where the reaction mediated by SPS is the limiting one (S. C. Huber et al., 1984; Ruan, 2014) (Fig 1A). We temperature adapted the measured maximum enzyme activity (*v* _max_) values by Arrhenius correction (see 2.5).

Generally, SPS *v* _max_ differed for controls and chilling night day 1 only in the night (06 h and 12 h), while chilling night day 7 showed higher SPS capacities besides of the 12 h time point (Fig. 2A, Tab. S2). In detail, for chilling night day 1 the SPS activities were similar to control, except of a reduction to a 0.3-fold level at the chilling night time points (06 h and 12 h). In contrast, chilling night day 7 showed higher SPS capacities which were on average 2.2-fold higher at day temperatures (22 *^◦^*C) compared to control, while at the end of the night (12 h) at 4 *^◦^*C levels were equal. Within the conditions, SPS *v* _max_ dropped in the night for all conditions compared to the day values and remained stable or similar during the light phase (Fig. 2A, Tab. S2).

**Figure 2:**
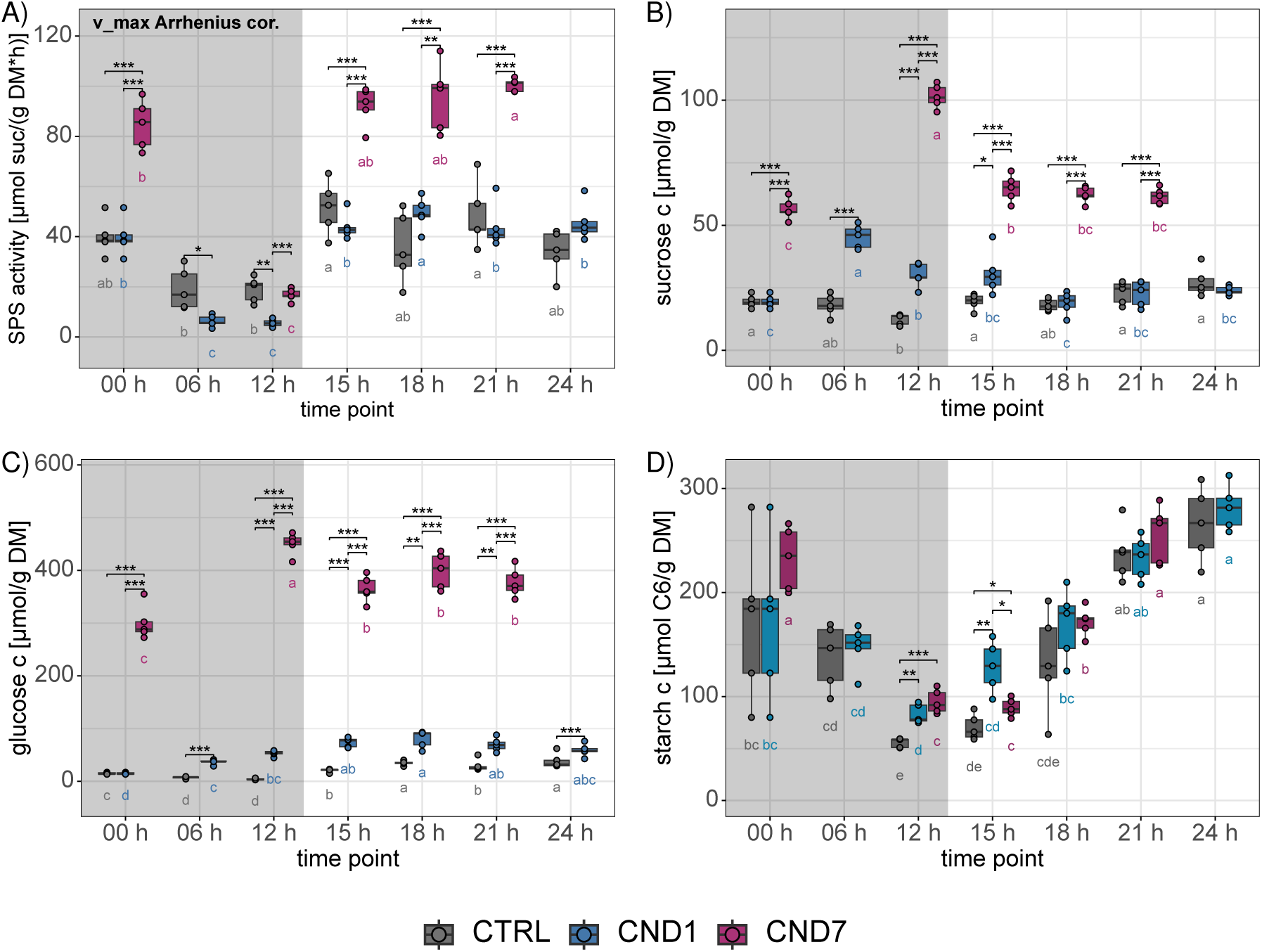
SPS *v* _max_ activity and sugar concentrations during time series of chilling nights (4 *^◦^*C/22 *^◦^*C day/night) events. **A)** Arrhenius corrected SPS *v* _max_ activities, **B)** sucrose concentrations, **C)** glucose concentrations, and **D)** starch concentrations in i) control (CTRL, gray, 18 *^◦^*C night and 22 *^◦^*C day), ii) chilling night day one (CND1, blue, 4 *^◦^*C night and 22 *^◦^*C day) and iii) chilling night day seven (CND7, magenta, 4 *^◦^*C nights and 22 *^◦^*C days). The carbohydrate concentrations in µmol per gram DM and *v* _max_ in µmol per gram DM per hour are plotted against the TPs [h]. The dark gray area indicates the nighttime, and light gray area the daytime. Five replicates were measured for each condition at each TP except for day seven 06 h and 24 h TP (no harvest). A t-test was performed between conditions at each TP (* p *<* 0.05, ** p *<* 0.01, *** p *<* 0.001). A one-way ANOVA with a Tukey HSD as a post hoc test (p *<* 0.05) was performed within each condition across TPs and is depicted as a CLD below each boxplot in the corresponding condition color. Abbreviations: ANOVA:= analysis of variance; c:= concentration; CLD:= compact letter display; CND1:= chilling night day 1; CND7:= chilling night day 7; CTRL:= control; DM:= dry mass; HSD=: honestly significant difference; SPS:= sucrose phosphate synthase.

The augmented neural ordinary differential equation (ANODE) model simulations showed an about 1.3-fold increase in SPS rates (*v* _pred_) for the second half of the day (18 h-24 h) of chilling night day 1 and an even higher about 2.6-fold increase during the seventh day, whereas both chilling night time points (06 h and 12 h) *v* _pred_ were decreased in chilling night day 1 to a 0.3-fold level (Fig. 3A, Suppl. Tab. S3, comparison between measured and modeled rates Fig. 4). At the beginning of the seventh chilling night SPS *v* _pred_ started at an 1.8-fold increased level, maintained it till the middle of the seventh chilling night (06 h), and then only decreased toward the end of the night to an 0.5-fold level of the control.

**Figure 3:**
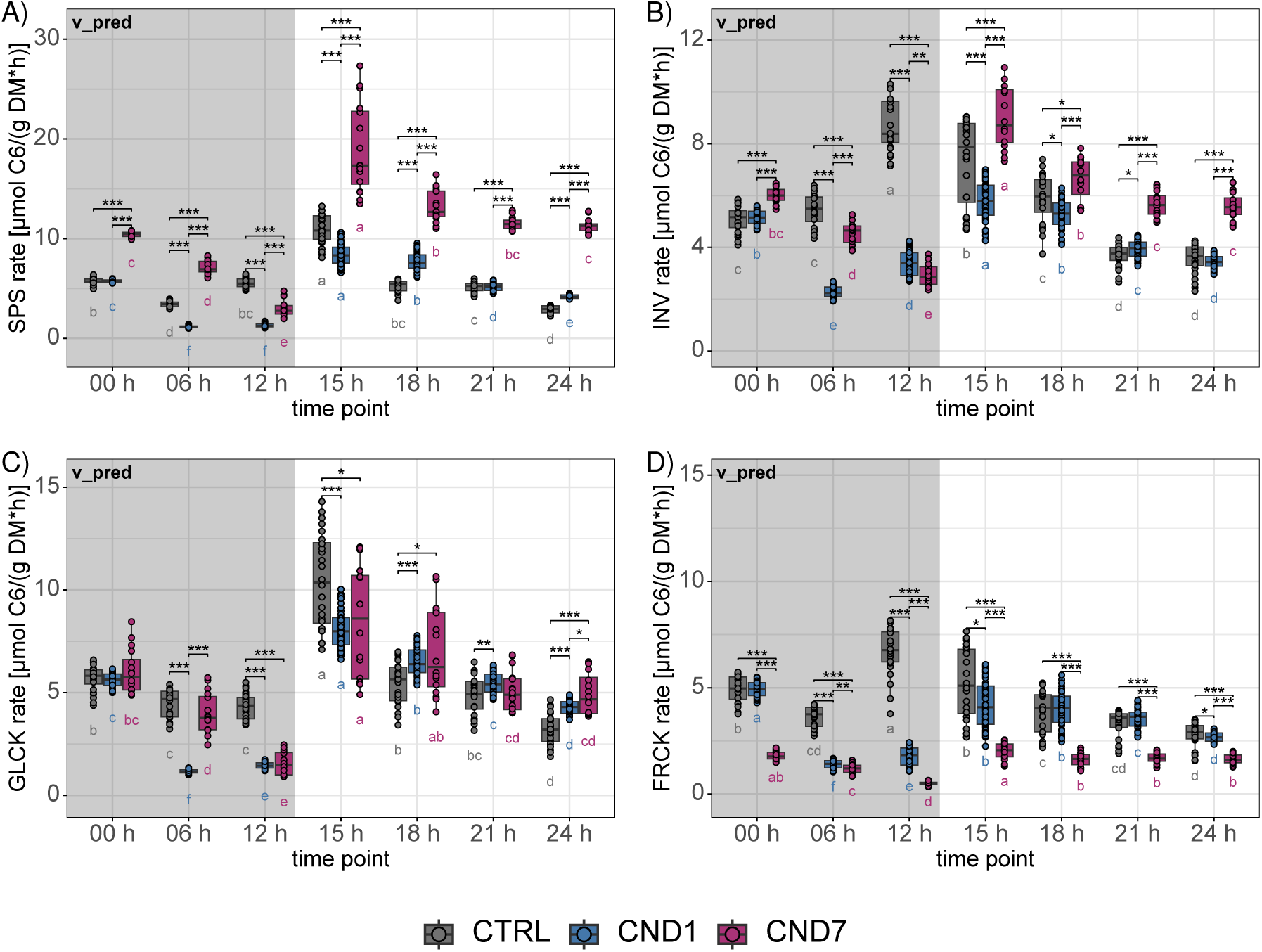
Predicted (*v* _pred_) ANODE modeled rates of key enzymes across chilling nights (4 *^◦^*C/22 *^◦^*C day/night). Rates were estimated by an ensemble of 100 augmented neural ordinary differential equation (ANODE) simulations per experimental condition (see. 2.7). Modeled activity rates of **A)** SPS (*v* _pred_), **B)** sum of INVs (*v* _pred_), **C)** GLCK (*v* _pred_), and **D)** FRCK (*v* _pred_) in i) control (CTRL, gray, 18 *^◦^*C night and 22 *^◦^*C day), ii) chilling night day one (CND1, blue, 4 *^◦^*C night and 22 *^◦^*C day) and iii) chilling night day seven (CND7, magenta, 4 *^◦^*C nights and 22 *^◦^*C days). The predicted rates *v* _pred_ in µmol C_6_ per gram DM per hour are plotted against the TPs [h]. The dark gray area indicates the nighttime, and light gray area the daytime. Predictions below the 0.2 and above the 0.8 quintile per condition and time point were removed for visual clarity. A t-test was performed between conditions at each TP (* p *<* 0.05, ** p *<* 0.01, *** p *<* 0.001). A one-way ANOVA with a Tukey HSD as a post hoc test (p *<* 0.05) was performed within each condition across TPs and is depicted as a CLD below each boxplot in the corresponding condition color. Abbreviations: ANODE:= augmented neural ordinary differential equation; ANOVA:= analysis of variance; CLD:= compact letter display; CND1:= chilling night day 1; CND7:= chilling night day 7; CTRL:= control; DM:= dry mass; FRCK:= fructokinase; GLCK:= glucokinase; HSD=: honestly significant difference; INV:=invertase; SPS:= sucrose phosphate synthase; TP:= time point.

**Figure 4:**
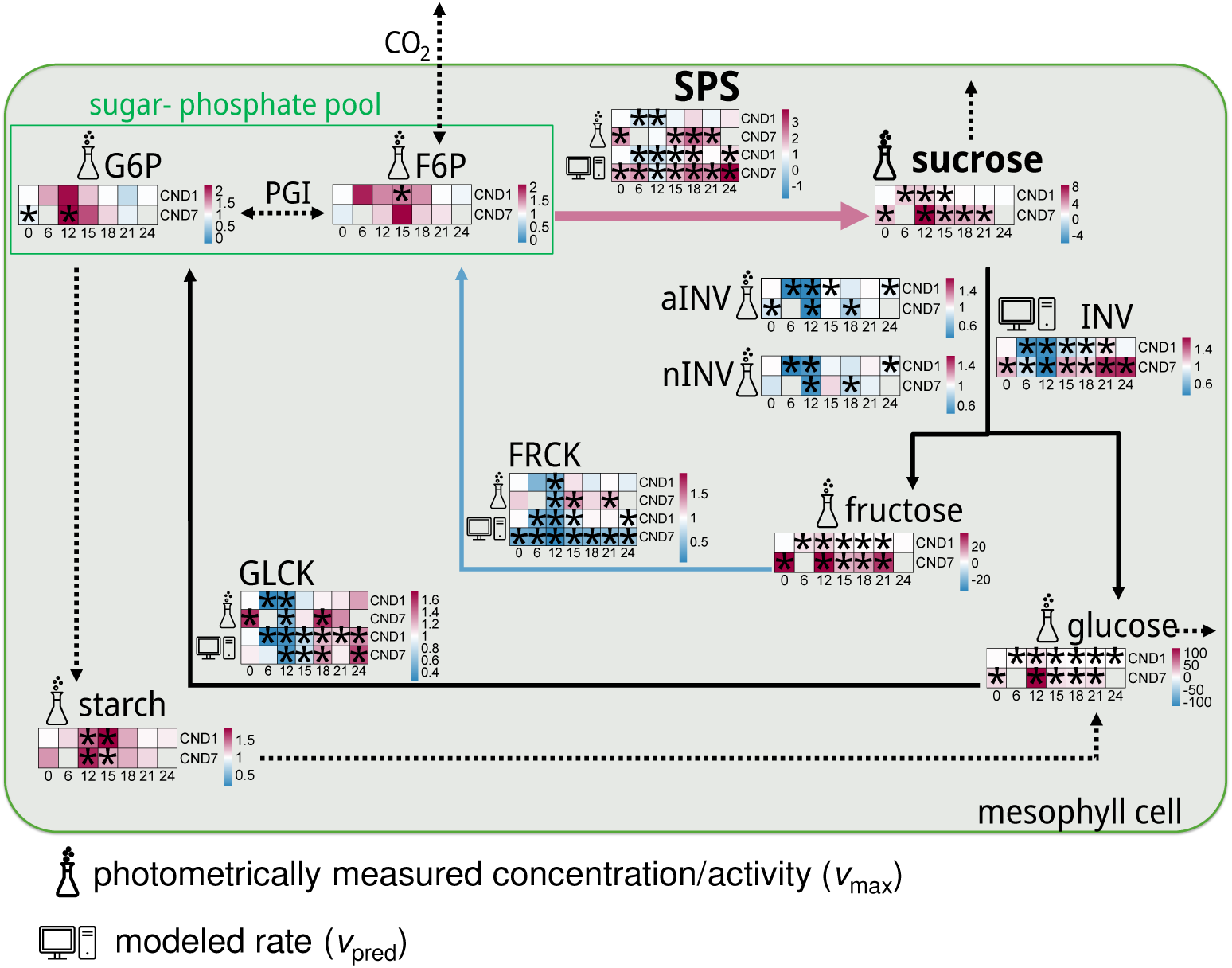
Carbon partitioning is shifted towards sucrose via increased SPS activity and lack of starch over-accumulation. Overview of metabolic and enzymatic changes under chilling nights normalized to CTRL embedded in a scheme of the carbohydrate metabolism. For each metabolite concentration, maximum enzyme activity *v* _max_ (Erlenmeyer flask), and modeled rate *v* _pred_ (computer icon) fold changes of chilling nights (4 *^◦^*C/22 *^◦^*C day/night), day 1 (CND1) and day 7 (CND7) normalized to control (CTRL) conditions are presented as heatmaps. Each heatmap has its own color legend, with high fold changes (FC*>*1) depicted in red and low fold changes (FC*<*1) in blue, stars indicate significant differences (p *<* 0.05, t-test). At the end of the light periods, no starch over-accumulations were observed for the chilling night samples while sucrose accumulated similar to permanent cold acclimation. Subsequently, analysis of enzyme activities via *v* _max_ (Erlenmeyer flask) and modeled rates *v* _pred_ (computer icon) showed a strong increase in SPS activities (red arrow), and a less pronounced increase in INV, and a by half reduced FRCK activity *v* _pred_ (blue arrow) in chilling night samples. Thus, the hypothesis arose that chilling nights led to a shift in carbon partitioning from starch to sucrose, opposing to what is known for permanent cold acclimation. Abbreviations: CND1:= chilling night day 1; CND7:= chilling night day 7; CTRL:= control; F6P:= fructose-6-phosphate; FC:= fold change; Frc:= fructose; FRCK:= fructokinase; G6P:= glucose-6-phosphate; Glc:= glucose; GLCK:= glucokinase; n/aINV:= neutral/acidic invertase; NPS:= net photosynthesis; Suc:= sucrose; SPS:= sucrose phosphate synthase.

When looking at F6P, a substrate of SPS, and the strong SPS activator G6P (Pattanayak, 1999; Giesbrecht et al., 2025), we determined slight increases in both chilling night day 1 and chilling night day 7 and a slow decrease during the day till the 21 h time point, where concentrations aligned to control conditions (Fig. 4, Suppl. Fig. S9A and S9B).

Overall, SPS *v* _max_ (maximum activity) and *v* _pred_ (modeled rates) were decreased significantly 0.3-fold in the cold hours of the first chilling night (4 *^◦^*C) and increased significantly 2.2-fold and 2.6-fold, respectively, during the seventh 22 *^◦^*C day (after six chilling nights) (Fig. 2A, 3A and 4, Suppl. Tab. S3).

### 3.4 Sucrose Accumulates Overnight under Chilling Night Conditions and Aligns with Controls on Day 1

Sucrose (suc) concentrations differed during the diurnal measurements, with all time points showing an about 3.1-fold increase in suc concentrations in chilling night day 7 compared to the control, while chilling night day 1 realigned with control during the day (Fig. 2B and 4, Suppl. Tab. S2 and S3). Specifically during the night, control samples reduced their suc amount, while chilling night samples increased their suc content (00 h vs 12 h). Chilling night day 1 samples showed a moderate accumulation of a 1.5-fold-change with a 2.5-fold increase at the middle of the night (06 h, Suppl. Tab. S3).

In contrast, chilling night day 7 started with a 2.9-fold higher content than controls at the beginning of the night (00 h), and nearly doubled its suc content over the course of the night. Within the first three hours of the day (15 h), different patterns were observed. Controls slightly increased the suc amount, chilling night day 1 remained stable and chilling night day 7 showed a rapid decrease to 0.6-fold of its initial level (Fig. 2B and 4, Suppl. Tab. S3). For all conditions, the suc level stabilized during the remainder of the day (Fig. 2B and 4, Suppl. Tab. S2).

Overall, chilling nights (4 *^◦^*C) induced temporary and long-term suc accumulation, with realignment of levels to control at midday in the first day, and permanent increase to about 3.1-fold level in chilling night day 7 (Fig. 2B and 4).

### 3.5 Elevated Hexose Levels and Distinct Hexokinase Responses after Chilling Nights

We further investigated sucrose cycling through suc cleavage by the invertases (INVs) (Fig. 1A) and their products glc and frc. Both acidic and neutral INVs showed a decreasing trend under chilling nights and following warm days in their *v* _max_ (Fig. 4, Suppl. Fig. S11A and S11B, Suppl. Tab. S3). This decrease was confirmed by our ANODE simulations at least for the first chilling night (*v* _pred_ decreased to 0.4-fold level) (Fig. 3B, Suppl. Tab. S4). In the following 22 *^◦^*C day INV rates *v* _pred_ realigned to control levels until midday (18 h). While INV *v* _pred_ was still decreased at the chilling night time points in chilling night day 7, during the day a 1.4-fold increase in INV rate was predicted in comparison to the control (Fig. 3B and 4, Suppl. Tab. S4).

The products of sucrose cleavage by the INVs, glucose and fructose, showed slightly different tendencies during the first day (Fig. 2C and 4, Suppl. Fig. S12, Suppl. Tab. S2). After the first chilling night, during the day, glc concentrations did not align to controls, which contrasted the findings of suc, whereas frc aligned to control within the first 24 h like suc (alignment at 18 h) but only at the end of the day (24 h) (2B and Fig. 2C, Suppl. Fig. S12). Overall, hexoses accumulated in chilling night samples compared to control conditions (Fig. 4, Suppl. Tab. S2). In detail, for glc, a 2.3-fold accumulation in chilling night day 1 and 11.5-fold in chilling night day 7 compared to controls was determined (midday, 18 h) (Fig. 2C and 4, Suppl. Tab. S3). Frc showed the same trend with a 1.4-fold increase in chilling night day 1 and a 14.0-fold accumulation on chilling night day 7 in the middle of the day (Suppl. Tab. S3 and Fig. S12).

For the specific diurnal trends, once again contrasting night results were found for the chilling night samples compared to controls. Glc decreased significantly (0.26-fold, Fig. 4, Suppl. Tab. S3) in controls and frc stayed stable, whereas both hexoses accumulated in chilling night plants, reaching up to 11.3-fold levels in chilling night day 1 for frc and 1.6-fold levels in chilling night day 7 for glc (00 h vs. 12 h).

For the following day, controls showed slight fluctuations during the light period for both hexoses. In contrast to suc, chilling night day 1 maintained constant glc levels equal to the levels observed at the end of the night, and frc realigned only at the end of the day (24 h) to the control levels. For chilling night day 7, glc and frc levels dropped in the first 3 h in the light (15 h), after which glc remained constant and frc showed another peak at the 18 h time point (Suppl. Tab. S2, Fig. 2C and Suppl. Fig. S12).

The two hexokinases GLCK and FRCK, which phosphorylate glc and frc, respectively, both tended to increase their daytime *v* _max_ towards chilling night day 7, while predicted rates (*v* _pred_) differed from these observations especially in FRCK. During the day time points of chilling night day 1, neither GLCK nor FRCK showed a significant difference in *v* _max_. On day 7, GLCK *v* _max_ fold change increased to a higher extent (1.6-fold, 18 h) than for FRCK, where an 1.4-fold increase was determined at 15 h (Fig. 4, Suppl. Fig. S11C and S11D, Suppl. Tab. S3). When considering the modeled rates, the first chilling night led to a decrease of *v* _pred_ GLCK rates (on average 0.5-fold) compared to the control, until the 18 h time point, at which rates realign (Fig. 3C, Suppl. Tab. S3). Already at the end of the day (24 h, day 1), a slight increase (1.3-fold) in GLCK *v* _pred_ was determined, which is also the average increase predicted for the second half of the seventh day (18 h-24 h).

For the first chilling night, FRCK *v* _pred_, on the other hand, is strongly reduced to 0.3-fold of the control condition, recovering towards control levels within the first three hour of the following 22 *^◦^*C day (15 h) (Fig. 3D and 4, Suppl. Tab. S4). On chilling night day 7, FRCK *v* _pred_ were on average decreased to a 0.4-fold level at all modeled time points. In parallel, glc and frc, the substrates for GLCK and FRCK, were both accumulated in plants under both chilling night day 1 and to an even higher extent (about 11.5-to 14.0-fold at the 18 h time point) in chilling night day 7, leading to a negative correlation between the substrates and their enzymes (Suppl. Fig. S13).

Overall, hexoses accumulated and negatively correlated with hexokinases towards chilling night day 7. Hexokinases themselves showed distinct patterns, with a reduction in GLCK rates (*v* _pred_) during chilling nights (4 *^◦^*C) with increases in the following 22 *^◦^*C, while FRCK rates (*v* _pred_) stayed continuously diminished throughout the seventh day (Fig. 2C, 3C, 3D and 4, Suppl. Fig. S13).

### 3.6 Seven Consecutive Chilling Nights do not Affect Starch Content

Starch concentrations were investigated as well, since starch is the primary energy storage of plants and a key component of carbon partitioning (Tiessen et al., 2002; Stitt et al., 2010; Thalmann & Santelia, 2017). Under all conditions, starch was degraded in the night and accumulated during the day (Fig. 2D and Suppl. Tab. S2).

Compared to controls, starch concentrations were only significantly higher at the 12 h time point (end of the night) and the 15 h time point (3 h of light) in both chilling night day 1 and chilling night day 7. The levels were 1.5- and 1.7-fold increased at 12 h time point and 1.8- and 1.3-fold increased at 15 h time point, respectively (Fig. 2D and 4, Suppl. Tab. S3). Additionally, in chilling night day 7 starch was decreased to a 0.69-fold level of chilling night day 1 at the 15 h time point. By midday (18 h), starch levels of chilling night samples realigned with control samples and continued the same accumulation trend until the end of the day (24 h) (Fig. 2D). Subsequently, no sustained starch over-accumulation upon chilling nights could be determined in *Arabidopsis* Col-0.

The course of net photosynthesis showed an increase in respiration during the night in chilling night day 1 as well as increasing means in photosynthetic activity in chilling night day 7 during the day (1.2-fold) (Suppl. Fig. S8A and Suppl. Tab. S2).

When looking at the starch to sucrose ratio, the increase in suc during the first chilling night led to an immediate decrease at the 06 h and 12 h time point compared to control (Suppl. Fig. S14B). On the following day after the first chilling night, the ratio tends to be slightly higher in the chilling night day 1 samples compared to control day time points (e.g. 18 h, midday). The accumulation of suc at chilling night day 7, and the fact that starch remained largely unaffected, led to lower starch/suc ratios (e.g. 1.3 at 18 h) compared to both control (3.7) and chilling night day 1 (4.5) (Suppl. Fig. S14).

In summary, nightly (4 *^◦^*C) starch degradation is decreased in the first chilling night, whereas during the first and seventh day (22 *^◦^*C) starch concentrations realigned to the control (Fig. 2D and 4).

## 4 Discussion

### 4.1 Chilling Nights

Fluctuating environmental temperatures force plants to employ highly specific acclimation mechanisms to maintain their physiological performance. In *Arabidopsis thaliana*, prolonged exposure to 4 *^◦^*C triggers a well-characterized cold acclimation program that reconfigures gene expression, regulatory networks, and carbohydrate metabolism, ultimately leading to the accumulation of starch and soluble sugars as energy reserves and osmoprotectants (C. L. Guy, 1990; Horton et al., 2016; Hoermiller et al., 2017; Adler et al., 2025). In natural habitats, however, annual plants are rarely exposed to prolonged cold; instead, they are more frequently exposed to “cool/chilling nights” - nocturnal temperatures of 0-6 *^◦^*C followed by daytime temperatures *≥*12 *^◦^*C higher (Suppl. Fig. S1). Such days with high temperature amplitudes most often occur in spring and autumn in which most ecotypes of *Arabidopsis thaliana* reach their vegetative phase (Meyerowitz, 1987) (Suppl. Fig. S1). This discrepancy raises important questions about how short-term, repeated cold events modulate the cold response pathways and which implications this has for the metabolic profile and enzyme activity of *Arabidopsis*. In this paper, we deciphered how *A. thaliana* Col-0 reacts to such chilling night conditions with nighttime temperatures of 4 *^◦^*C followed by moderate daytime temperatures of 22 *^◦^*C at day 1 and day 7. Based on a comprehensive time series analysis of chilling nights, we identified fluctuations in carbohydrate content and the associated enzymes that differ from those observed under well described permanent cold conditions and appear to be unique to chilling nights.

### 4.2 Biologically Constrained ANODE Modeling Reveals Plausible Reaction-Rate Dynamics

Standard ANODE models (Dupont et al., 2019) risk learning system dynamics in abstract, non-physiological spaces, where recovered quantities may not correspond to real reaction rates. While such models excel in predictive accuracy, their lack of interpretability limits their utility for mechanistic inference. By fixing the stoichiometric structure of the metabolic network a priori and learning only the fluxes through this network, it was ensured that the estimated fluxes were physiologically meaningful in magnitude. However, the functional dependence of each flux on metabolite concentrations remains learned rather than mechanistic; thus, while the recovered flux magnitudes are biologically interpretable, the underlying kinetics and regulatory mechanisms are not explicitly resolved. This distinction is critical: this approach yields interpretable flux values (*v* _pred_) but not interpretable kinetic laws.

The non-identifiability arising from sparse time-series data is a fundamental challenge in flux estimation. While bounding reaction rates by their *v* _max_ values constrained the solution space to biologically plausible regimes, it did not eliminate non-identifiability entirely. The ensemble approach addresses this by quantifying uncertainty in the inferred fluxes, but it is important to note that this uncertainty is conditional: it reflects variability across fits that share the same stoichiometric network and model architecture. It does not account for: miss-specification of the reaction network (e.g., omitted or incorrect reactions), architectural biases (e.g., the choice of neural network architecture), systematic errors in *v* _max_ measurements, or unmodeled regulatory mechanisms (e.g., allosteric control).

Thus, while ensemble variability provides a lower bound on uncertainty, additional sources of error may exist. The obtained results should therefore be interpreted as hypotheses about plausible flux distributions under the assumed network topology, with differences between conditions classified as either robust (consistent across ensemble members) or tentative (falling within ensemble variability). Future work could systematically vary the network stoichiometry and model architecture to assess the sensitivity of these conclusions, though such efforts would require substantially more data and computational resources.

This work demonstrates that constraining ANODE models with biological priors (e.g., fixed stoichiometry and enzyme activities as *v* _max_) can bridge the gap between predictive performance and mechanistic interpretability. However, the trade-off between interpretability and mechanistic fidelity remains: while biologically plausible flux magnitudes were recovered, the dynamics that produce them remain opaque. Addressing this will likely require integrating time-resolved metabolomics, enzyme activity assays, and targeted perturbations to disambiguate the learned relationships between metabolites and fluxes.

### 4.3 Chilling Nights Enhance Nocturnal Sucrose Accumulation and SPS Activity

The concentration of sucrose increased 2.5-fold upon the first chilling night (immediate stress response), peaked at the middle of the night (06 h) and decreased again to control level at midday (18 h) (Fig. 2B and 4, Suppl. Tab. S2 and S3). After seven chilling nights, suc levels were increased about 3-fold (Fig. 2B), which is within the range of suc accumulation reported for acclimation to permanent cold (Kitashova et al., 2021, 2023).

The accumulation of suc in the first chilling night could not be explained by the measured *v* _max_ of SPS or INV as they were largely unaffected (Fig. 2A and Suppl. Fig. S11A and S11B). In the model results, in the first chilling night SPS *v* _pred_ rates were significantly decreased on both night time points (06 h and 12 h) to an about 0.3-fold level of the control (Fig. 3A). Therefore, it seems most likely that the increase of suc in the first half of the first chilling night might be explained by either a reduction of suc export to the sink organs (Krapp & Stitt, 1995; Å. Strand et al., 1997), by a reduction in cellular respiration (Huner et al., 1998) or alteration of alternative pathways (e.g., cleavage of suc by sucrose synthase (SUS)).

We determined elevated nightly respiration rates via CO_2_-gas exchange measurements (net photosynthesis) in the middle of the night (6 h) with a fold change of 1.7-fold (*±* 1.4, not significant due to high variance of biological replicates) compared to controls (Suppl. Fig. S8A) pointing towards a reduced export to the sink and/or other reactions.

In acclimation to permanent cold, accumulation of suc is reported due to a retardation in plant growth, which outweighs the reduced photosynthetic activities reported there (Huner et al., 1998; Fürtauer et al., 2019). Since suc biosynthesis is the partitioning pathway for sugar phosphates besides starch biosynthesis, the accumulation of suc on the seventh day of chilling nights might also explain the lack of starch accumulation (Fig. 2D and Suppl. Tab. S2) in comparison to acclimation to permanent cold (Kaplan & Guy, 2004; Hoermiller et al., 2017; Kitashova et al., 2021, 2023).

An increased suc synthesis may simply lead to an educt depletion for starch biosynthesis. The increase in suc biosynthesis is supported by the 2.2-fold increase in *v* _max_ of SPS and 2.6-fold increase in modeled enzyme rate *v* _pred_ on the seventh day as well (Fig. 2A and 3A).

A positive correlation between starch and suc concentrations was found in the control samples, while increasingly negative correlations were found in the first and seventh chilling night (Suppl. Fig. S13). This change of correlations in chilling nights indicates, that the flux from shared substrates (i.e. sugar phosphates) increased towards sucrose and was maintained for starch. F6P, which is substrate for suc synthesis, and G6P, which is an activator of SPS (Pattanayak, 1999; Giesbrecht et al., 2025), are both quite stable in their diurnal rhythm (Suppl. Fig. S9A and S9B). Both are, on average, 1.3-fold increased under chilling night conditions (Suppl. Tab. S3) and could thereby contribute to increased suc synthesis. This can also be seen in the positive correlation of suc with both G6P and F6P under chilling night conditions (Suppl. Fig. S13B and S13C).

Generally, the activity of SPS is an important regulator in the central carbohydrate metabolism because it is limiting for the export of triose phosphates from the chloroplast to the cytosol. When triose phosphates accumulate in the chloroplast, a disturbance of the photosynthetic equilibrium can occur, leading to formation of reactive oxygen species, which in turn can, at a certain extent, cause cell death (Poolman et al., 2000; Schneider et al., 2002; Nägele et al., 2012; Sharma et al., 2012). Thus, the tolerance towards cold temperatures can be enhanced due to increased SPS capacity in *Arabidopsis thaliana* as shown in SPS over expression lines by Å. Strand et al. (2003) and Herrmann et al. (2019). Noticeably, in acclimation to permanent cold no significant increase of the SPS *v* _max_ was measured after seven days at 4 *^◦^*C (Kitashova et al., 2021, 2023), which contrasts the here described findings (Fig. 2A and 3A). This difference might occur because the need for an increased triose phosphate export on the first hand arises, through rapid changes in the conditions (Nägele et al., 2012), like the temperature amplitude applied here.

Overall, acclimation after seven chilling nights led to sucrose accumulation and increase in both SPS *v* _max_ and modeled *v* _pred_ SPS rates in Col-0 *Arabidopsis thaliana* (Fig. 4). We conclude, that there is a shift towards increased sucrose biosynthesis.

### 4.4 Chilling-induced Hexose Accumulation is Linked to Lower HXK Rates

The concentrations of the hexoses, glucose and fructose increased upon the immediate stress response to the first chilling night, peaking at different time points, and starting to decrease again within the first 24 h (Fig. 2C, Suppl. Fig. S12, and Suppl. Tab. S2).

After acclimation to seven consecutive chilling nights, glucose was increased about 11.5-fold (18 h, midday) (Fig. 2C and 4), which is within the range of accumulation reported for both glc and frc under acclimation to permanent cold (Kitashova et al., 2021, 2023). Glucose plays a crucial role in signaling during cold stress, particularly for glc-sensing enzymes like suc non-fermenting-related kinase 1 (SnRK1), hexokinase, and target of rapamycin (TOR). These enzymes can activate various transcription programs, thereby influencing the expression of specific cold-related genes (Rolland et al., 2006; Baena-González & Sheen, 2008). Additionally, glc is involved in the abscisic acid (ABA) pathway, which increases the level of endogenous ABA and promotes the expression of genes associated with ABA synthesis and signaling (Cheng et al., 2002). Both ABA and glc are capable of inducing the expression of essential regulators in response to abiotic stress, such as CBF3 (C-repeated binding factor), COR15A (cold-regulated), and RD29A (responsive to dehydration) (Yuanyuan et al., 2009).

Fructose, on the other hand, was accumulated to an even higher extent (14.0-fold, 18 h midday) in acclimation to chilling nights (chilling night day 7, Suppl. Fig. S12 and Suppl. Tab. S2) than reported in acclimation to permanent cold (10-to 11-fold (Kitashova et al., 2021, 2023)). This might indicate an increased need of frc in signaling pathways e.g. as a signal for phosphorylated sugars such as fructose 2,6-bisphosphate, which is considered a controlling unit for partitioning of photoassimilates and sucrose synthesis (McCormick & Kruger, 2015; Scharte et al., 2023), or in the regulation of, for example, SPS (Giesbrecht et al., 2025).

In theory the accumulation of glc and frc could be explained by increasing INVs activities for which no indication could be found in the measured *v* _max_ values of either the neutral or acidic INV. The *v* _max_ of both acidic and neutral invertase were decreased on the seventh cold night and especially the following warm day (Suppl. Fig. S11 and Suppl. Tab. S2). Moreover, model simulations predicted that INV *v* _pred_ decreased significantly to 0.8-fold at the 15 h time point of chilling night day 1, whereas it increased significantly by 1.4-fold on average at the day time points on the seventh day (Fig. 4, Suppl. Tab. S4). Therefore, except for the seventh day, INV *v* _pred_ cannot explain the observed accumulation of both glc and frc throughout chilling nights.

As the two INVs, of which maximum activities were measured here, are located in different compartments of the cell, the neutral in the cytosol and the acidic (among others) in the vacuole, an in-depth subcellular analysis of the metabolite concentration would be of interest, since it was shown that i.e. vacuolar (acidic) invertase stabilizes photosynthesis under cold stress (Weiszmann et al., 2018). The maximum activity *v* _max_ of the acidic invertase was generally higher and the decrease in the fold changes slightly stronger than for the neutral invertase (Suppl. Fig. S11A and S11B). Therefore, the level of decrease is overall way stronger in the acidic invertase. This might hint at compartment specific regulations to enable accumulation of osmotically active and cryoprotective substances according to the specific needs of the different compartments (Gerhardt & Heldt, 1984; Fürtauer & Nägele, 2016; Fürtauer et al., 2016; Hoermiller et al., 2017).

In *Arabidopsis*, a shift of suc cleavage from the vacuole to the cytosol takes place in cold-susceptible accessions like C24, while cold-tolerant ecotypes like Rsch showed the opposite trend when exposed to permanent cold (Nägele & Heyer, 2013; Weiszmann et al., 2018). In acclimation of Col-0 to permanent cold (4 *^◦^*C) either no significant difference (Kitashova et al., 2023) or a stronger decrease to 0.4-fold levels were reported for both *v* _max_ of acidic and neutral invertases (Kitashova et al., 2021). Therefore, the change in *v* _max_ invertase activities under chilling nights in Col-0 are within the margins found under acclimation to permanent cold.

In the first night (00 h till 12 h), no increase in *v* _max_ of INVs (Fig. 3B, Suppl. Fig. S11A and S11B) or SPS (Fig. 2A, 3A) could be measured or estimated by modeling (*v* _pred_), a decrease in hexokinase activities could also be responsible for the accumulation of glc and frc (Fig. 4). While *v* _max_ of FRCK were about 1.2-fold increased on the seventh day, during the night FRCK *v* _max_ was 0.4-fold decreased under both chilling night day 1 and chilling night day 7 (Suppl. Fig. S11 and Suppl. Tab. S2). This decrease of FRCK *v* _max_ during the night could at least partially account for the accumulation of frc during chilling nights. This was additionally confirmed by FRCK modeled rates *v* _pred_, which not only decreased to 0.3-to 0.4-fold of control during the first chilling night (06 h and 12 h) and further down to 0.1-fold at the end of the seventh chilling night (12 h), but also stayed reduced to 0.5-fold of the control *v* _pred_ throughout the seventh day (Fig. 3D and Suppl. Tab. S4). This influence of decreased FRCK *v* _pred_ on frc accumulation can also be seen in the negative correlation between both, which arises already upon the first chilling night (Suppl. Fig. S13). Under acclimation to permanent cold, no significant change of FRCK *v* _max_ were reported (Kitashova et al., 2021, 2023), matching also our measured *v* _max_ values at the 18 h time point (Suppl. Fig. S11D).

The discrepancy of glc and frc accumulation, could also arise from sucrose synthase (SUS) activities, which can split suc into frc and UDP-glc or ADP-glc instead of glc (Bolouri-Moghaddam et al., 2010). Therefore, it would be interesting to see if SUS enzyme activities behave similar to INVs and if the implementation of SUS can further shed light into these imbalanced observations (Bolouri-Moghaddam et al., 2010). Further, the lack of SUS implementation could have also led to an overestimation of the FRCK activity reduction by *v* _pred_.

When considering *v* _max_ of GLCK, a slightly stronger impact of chilling nights was measured, since GLCK *v* _max_ increased up to 1.6-fold on day 7 (Suppl. Fig. S11C), which is slightly lower than the 2.0-fold increase in GLCK *v* _max_ reported for acclimation to permanent cold at midday (Suppl. Tab. S2 comparison to (Kitashova et al., 2021, 2023)). Like with FRCK before, the decrease of GLCK *v* _max_ in the night (∼0.3-fold on day 1 and ∼0.5-fold on day 7) could contribute to glc accumulation.

It was shown previously via a kinetic modeling approach that hexokinases play a crucial role in the stabilization of suc cycling when plants experience abiotic stressors (Nägele et al., 2010; Henkel et al., 2011). In our model simulations, the GLCK rate *v* _pred_ was decreased to a 0.3-fold level in the first chilling night as well, while in the seventh chilling night only for the 12 h time point a decrease to a 0.4-fold level was predicted (Fig. 3C, Suppl. Tab. S4). The increase in both FRCK and GLCK *v* _max_, which was at least for GLCK on the seventh day confirmed by the modeled rates *v* _pred_, could have led to the assumption of a higher need for G6P and F6P, which was not supported by the very slight average increase of ∼1.3-fold in G6P and F6P concentrations and the strong decrease in modeled FRCK rate *v* _pred_ (Suppl. Tab. S2, Fig. 3C and 3D Suppl. Fig. S3).

The low increase of the G6P concentration is of special significance because it was shown that the formation of G6P via GLCK is vital for plant growth and fitness, as many biosynthetic pathways rely on this step (Vanderwall & Gendron, 2023). In cold acclimated plants, G6P concentrations are increased 3-fold (Kitashova et al., 2023), under chilling nights at midday of the seventh day (18 h) only a non significant increase of 1.2-fold was determined (Suppl. Fig. S9B). This indicates that either under chilling night conditions there is less need for accelerated growth than under acclimation to permanent cold or the normal daytime temperature is sufficient to maintain standard growth, as the rosettes grow and gain weight (Suppl. Fig. S16).

Taken together, while accumulation of glc was unsatisfactory explained by the analyzed variables of the carbohydrate metabolism in this study, the high accumulation of frc might be explained to some extent by the decrease in the modeled rate *v* _pred_ of FRCK, which could not have been derived from the measured *v* _max_ values alone.

### 4.5 Chilling Nights Do Not Induce Starch Over-Accumulation

Typically, under permanent cold a 2-to 10-fold accumulation of starch has been reported for many *Arabidopsis* ecotypes, including Col-0 (Suppl. Tab. S3) (Hannah et al., 2006; C. Guy et al., 2007; Nagler et al., 2015; Kitashova et al., 2021, 2023). Contrary to permanent cold acclimation, several chilling nights did not led to a sustained over-accumulation of starch (Fig. 2D). Although starch levels were higher at the end of the night under chilling-night conditions (1.5-fold chilling night day 1 and 1.7-fold chilling night day7), this increase was transient and did not persist into a strong daytime accumulation (n.s. from midday (1.3-fold) onward for both chilling night day 1 and 7) (Fig. 4, Suppl. Tab. S3). Detailed, at the end of the night, starch levels were significantly higher for the chilling night conditions (12 h), besides of starting at similar levels (high variance at control). Calculating the overall nightly degradation rate of starch, showed two distinct pictures. While starch degradation rates (00 h till 12 h) decreased for the first chilling night (∼-7 µmol·g^-1^ DM·h^-1^), it was even slightly increased for the seventh chilling night(∼-11 µmol·g^-1^ DM·h^-1^) compared to control (∼-10 µmol·g^-1^ DM·h^-1^) conditions (Suppl. Tab. S14). Consecutively, within the first three hours of the day (12 h till 15 h), chilling night samples showed higher starch levels compared to controls. Within all conditions no significant starch accumulation was observed within these hours, but means increased for controls and chilling night day 1 and slightly decreased for chilling night day 7.

The decline of the degradation rate in the first cold night might be due to thermodynamic effects that lead to a reduced activity in starch degrading enzymes. The rules of Van’t Hoff and Arrhenius state, if simplified, that with an increase or decrease of the temperature over about 10 *^◦^*C, enzyme reaction rates increase or decrease about 2 to 3-fold, respectively (Van’t Hoff, 1884; Johnson & Thornley, 1985). The product of starch degradation by *β*-amylase, which hydrolyses *α*-1,4-D glucosidic bonds in polysaccharides, is maltose, which serves as an osmo-protectant and as substrate for hexoses, raffinose, and proline biosynthesis (Kaplan & Guy, 2004; Sicher, 2011; Ribeiro et al., 2022). Both the fact that starch degradation is up-regulated through an increase in the *β*-amylase activity to increase the amount of cryoprotectants in cold acclimation and the short amount of time the plants were exposed to the chilling temperature at this time point, lead to the assumption that the decreased degradation rate is not caused by cellular regulation (Kaplan & Guy, 2004; Sicher, 2011; Ribeiro et al., 2022).

On the other hand, there are reports which show a down-regulation in the *α*-amylase activity in rice plants under cold stress (W. Wang et al., 2016). It is also known that the different amylases (*α*-, *β*-and iso-amylase) can be regulated and expressed differently in various genotypes of *A. thaliana*, which means that a cellular regulation can not be excluded completely (Nagler et al., 2015). Furthermore, we can not elucidate from starch concentrations alone whether an increase in starch degradation rates occurs also during the light phase, which could have also led to the observed lack of over-accumulation. Therefore, measurements of either the amylase activities or maltose concentrations would be helpful to provide insights in this multi-factorial system. Especially as maltose is feeding into the glc pool as well, which is why decreasing maltose concentrations could contribute to the measured glc accumulation.

A third possibility, according to the rule of Le Chatelier, might be a shift in the metabolic equilibrium, which could occur due to reduced cellular respiration, resulting in an accumulation of the sugars obtained by starch breakdown (Treptow, 1980). The measured net photosynthesis values at that time point are contradictory to this hypothesis. There is even a significant increase in the measured respiration at the end of the first chilling night at the 12 h time point in comparison to control (Suppl. Tab. S2 and Suppl. Fig. S8A). The delayed increase of starch concentration on day seven by about three hours might be caused by a combination of two of the above described effects. If, after six cold nights, the *β*-amylase activity was increased, as reported for acclimation to permanent cold (Kaplan & Guy, 2004; Ribeiro et al., 2022), the low temperatures might lead to a prolongation of the period at the beginning of the day during which the *β*-amylase activity is down-regulated again, allowing for starch accumulation. Overall, the absence of significant starch over-accumulation under chilling night conditions indicates that alternating chilling nights and warm days elicit a different carbon partitioning response than permanent cold acclimation, where starch accumulation is much stronger. Thus, it seems that when each cold night is followed by a warm day, plants do not prepare for freezing in the manner observed in permanent cold where starch is accumulated to protect the plant from freezing and to supply energy as well as cryoprotective sugars (Dong & Beckles, 2019). The increase in SPS rate might reduce the need for starch accumulation, increasing the freezing tolerance by means of increasing photosynthesis activity instead, since increasing freezing tolerance was reported in SPS over expression lines (Å. Strand et al., 2003; Herrmann et al., 2019). Here, in contrast, the starch/sucrose ratio showed a clear shift toward sucrose in chilling night day 7 (1.2-1.4, midday) (Suppl. Fig. S14B) and no starch over-accumulation like in permanent cold (2.8-4.9, midday (Kitashova et al., 2021, 2023)) was observed. The reduction of the starch/suc ratio to 0.4-fold relative to the control suggests that carbon partitioning and energy storage under chilling-night conditions are shifted toward sucrose rather than starch, which contrasts with the response observed during acclimation to permanent cold.

### 4.6 Carbon Partitioning is Shifted Towards Sucrose Under Acclimation to Chilling Nights in *Arabidopsis*

Since climate change leads to more rapid changes in temperatures and increases in the amplitude between minimum and maximum temperatures throughout the year (Suppl. Fig. S1), plants need to adapt to these conditions. We showed in this study that, similar as in acclimation to permanent cold, Col-0 *A. thaliana* shows a clear distinction between the immediate stress response upon the first chilling night and acclimation to seven consecutive chilling nights (Fig. 1B, correlations Suppl. Fig. S13B and S13C).

In the immediate response to the first chilling night, soluble sugars were increased during the chilling night but realigned with control levels within the following warm day, except for glucose which remains elevated even towards the end of the day (Fig. 4) highlighting its importance as signaling molecule under abiotic stress (Cheng et al., 2002; Rolland et al., 2006; Baena-González & Sheen, 2008; Yuanyuan et al., 2009). On the following warm day, *v* _max_ enzyme activities were largely unaffected by the first chilling night. Even though, the modeled simulations (*v* _pred_) indicated decreased enzyme rates during the chilling night (06 h and 12 h), to various degrees for SPS, INV, GLCK, and FRCK, as well as slightly increased SPS rates (*v* _pred_) during the following warm day (15 h till 24 h) (Fig. 4, Suppl. Tab. S4). The different diurnal trends and their different adaptation to the chilling night and the following warm day, show the importance of time series analysis.

We could show that acclimation to prolonged chilling night events (chilling night day 7) differs clearly from acclimation to permanent cold. We determined an accumulation of soluble sugars such as sucrose, glucose and fructose. Compared with previous reports on permanent cold (Kitashova et al., 2021, 2023), fructose levels appeared higher, whereas sucrose and glucose increased to similar extents. Chilling night conditions led to a significant 2.2-fold increased of SPS *v* _max_, whereas SPS *v* _max_ was unchanged (non-significant increase) in permanent cold (Kitashova et al., 2021, 2023). Our ANODE modeling approach based on the measured metabolite concentrations and maximum enzyme activities, could further back up this hypothesis by a predicted SPS *v* _pred_ rate that is 2.5-fold increased compared to the control (18 h). Even though maximum INV activities (*v* _max_) tended to be decreased, the modeled approach showed an increase in INV *v* _pred_ and GLCK *v* _pred_ on the seventh day (15 h to 24 h). Solely, FRCK remains at a decreased *v* _pred_ rate compared to the control, hinting that FRCK might play an important role in the acclimation to chilling nights and an increased need for frc which is with a 14.0-fold increase even higher accumulated than under permanent cold (10-to 11-fold) (Kitashova et al., 2021, 2023). The increased SPS rate along with the sucrose accumulation, while starch levels remain stable throughout the acclimation to chilling nights, hints at crucial difference in carbon partitioning and energy management of Col-0 plants experiencing chilling night events in comparison to permanent cold, since starch is known to accumulate under permanent cold up to 10-fold (Kitashova et al., 2021, 2023). Even though suc and starch accumulation as well as decreased photosynthetic activity in prolonged permanent cold are attributed to a retardation in plant growth (Huner et al., 1998; Fürtauer et al., 2019; Ribeiro et al., 2022), this was not the case for suc accumulation under chilling nights conditions, since the rate of plant growth was not reduced (Suppl. Fig. S16), photosynthesis activity was even slightly elevated on the seventh day and starch did not accumulate in the first place. Confirming the advantage of higher SPS activities by e.g. SPSA1 mutants and/or over-expression lines and their impact on chilling night carbohydrate partitioning and freezing tolerance together with measurements of starch degrading enzymes would be of interest for further investigations. Additionally, consideration of more cold-sensitive and cold-tolerant natural *A. thaliana* accessions besides Col-0 might help to elucidate adaptation strategies.

Overall, our results demonstrate that chilling nights induce a distinct acclimation response in *A. thaliana* (Col-0) compared with permanent cold. This response is characterized by a shift in carbon partitioning toward sucrose biosynthesis, driven by elevated SPS rates (measured and modeled), while starch over-accumulation is absent.

## Acknowledgments

We want to thank members of the Fürtauer, van Dongen and Matuszyńska groups for valuable input into this project, as well as Anja Reinstädler, Agnieszka Miler-Bartkowiak and Brigitta Ehrt for technical support. We also thank Pia Falter, Kira Neumann and Lisa Reinmuth for harvesting support, and Joshua Bonn for sharing coding advice.

## Competing Interest

The authors declare no conflicts of interest.

## Author Contribution

L.F. designed the experiments. C.B. performed experiments with the help of O.G., M.B. and S.J.. M.v.A. developed the model and performed numerical simulations with the guidance of A.M.. C.B. and A.J. computationally analyzed the data. C.B. and L.F. evaluated the data. All authors discussed the results critically. C.B. wrote the main draft with input of A.J., M.v.A. and L.F.; J.T.v.D. and A.M. revised the manuscript.

## Data Availability

The data that support the findings of this study are available in the supplementary material of this article. Source code and data required to reproduce the ANODE analyses are available at: https://github.com/Computational-Biology-Aachen/ChillSugar.

## Funding

C.B. was fully and A.J. was partially funded by L.F.s WISNA program, A.J. additionally via the CLAR project (Stichting Jan IngenHousz, Jan IngenHousz Institute Wageningen).

## A Supporting Information

### A.1 Chilling Nights

**Figure S1:**
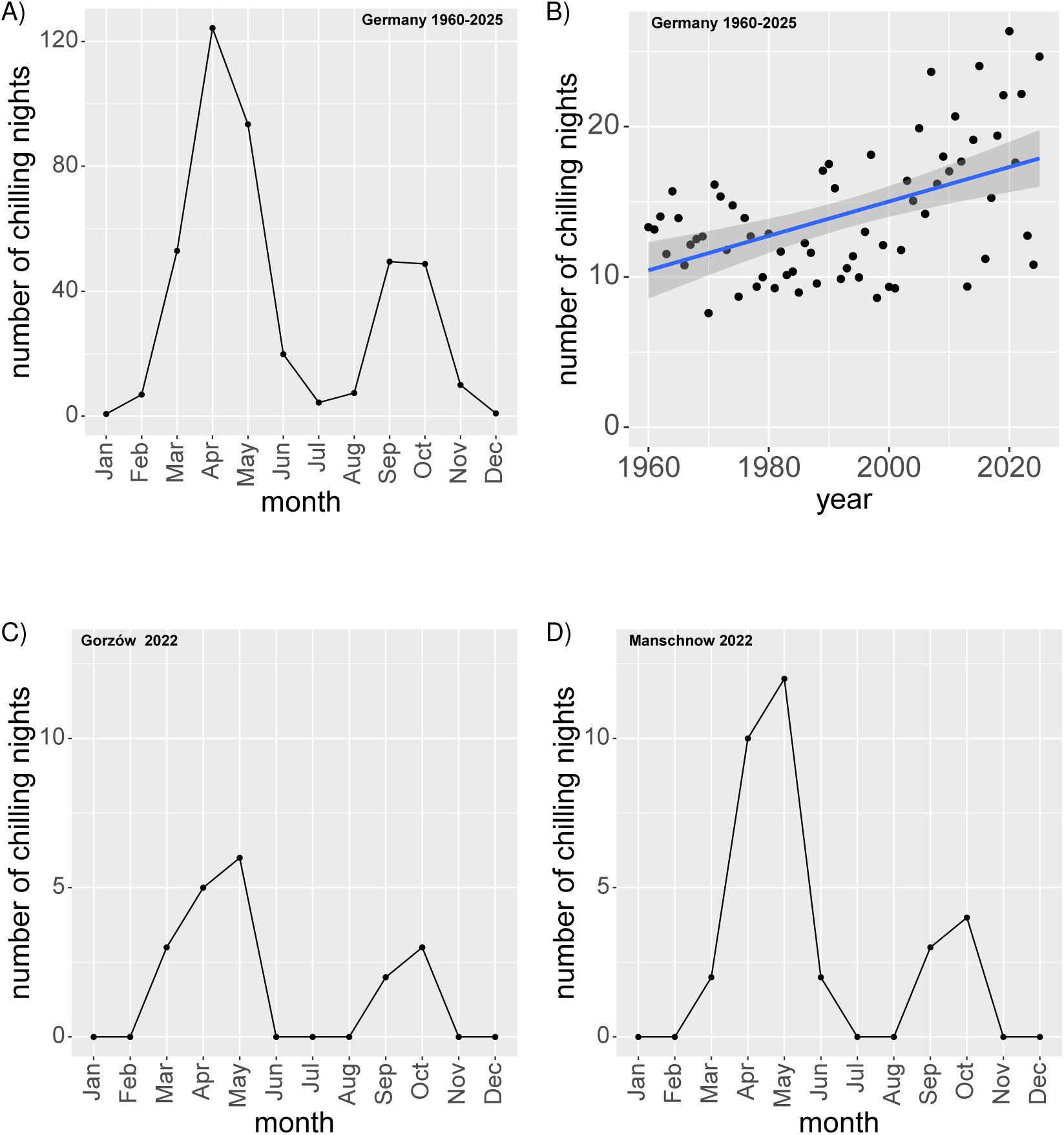
Chilling night events. **A)** The mean number of chilling night events per month that occurred in Germany throughout the years from 1960 to 2025 per active weather station (>= 100) throughout the years (Wetterdienst, 2026). **B)** The number of chilling night events in Germany are depicted per year normalized to the number of active weather station in each year (Wetterdienst, 2026). The blue line shows the linear correlation of the data, with a 95% confidence interval depicted in dark gray. **C)** Chilling Nights in Gorzów (Poland) 2022. **D)** Chilling Nights in Manschnow 2022 (Germany), closest weather station to Gorzów within Germany.

### A.2 Supplemental Material and Methods

#### A.2.1 Metabolite Analysis

##### Sucrose, Glucose and Fructose Quantification

The soluble sugars (non-phosphorylated) were extracted from dried leaf material by double ethanolic extraction (80% ethanol) at 80 *^◦^*C shaking at 350 rpm for 30 min. After centrifugation (21,100 g for 10 min) supernatants were collected in a new reaction tube. The supernatant was dried in a vacuum concentrator and sugars resolved in ddH_2_O shaking at 750 rpm for up to 180 minutes. The resulting plant extract was centrifuged and the supernatant was transferred into a clean reaction tube.

**Sucrose** was quantified using a modified version of an anthrone assay described by Dreywood (1946). For the quantification, plant extract was mixed with a 30% KOH solution and incubated at 95 *^◦^*C for 10 min. Subsequently, the samples were cooled down to room temperature. Anthrone reagent (0.14% (w/v) of anthrone in 14.6 M H_2_SO_4_) was added to the sample. Through incubation at 40 *^◦^*C for 30 min, a bluish-green complex is formed, which was photometrically determined at 620 nm (Suppl. Fig. S2).

**Figure S2:**
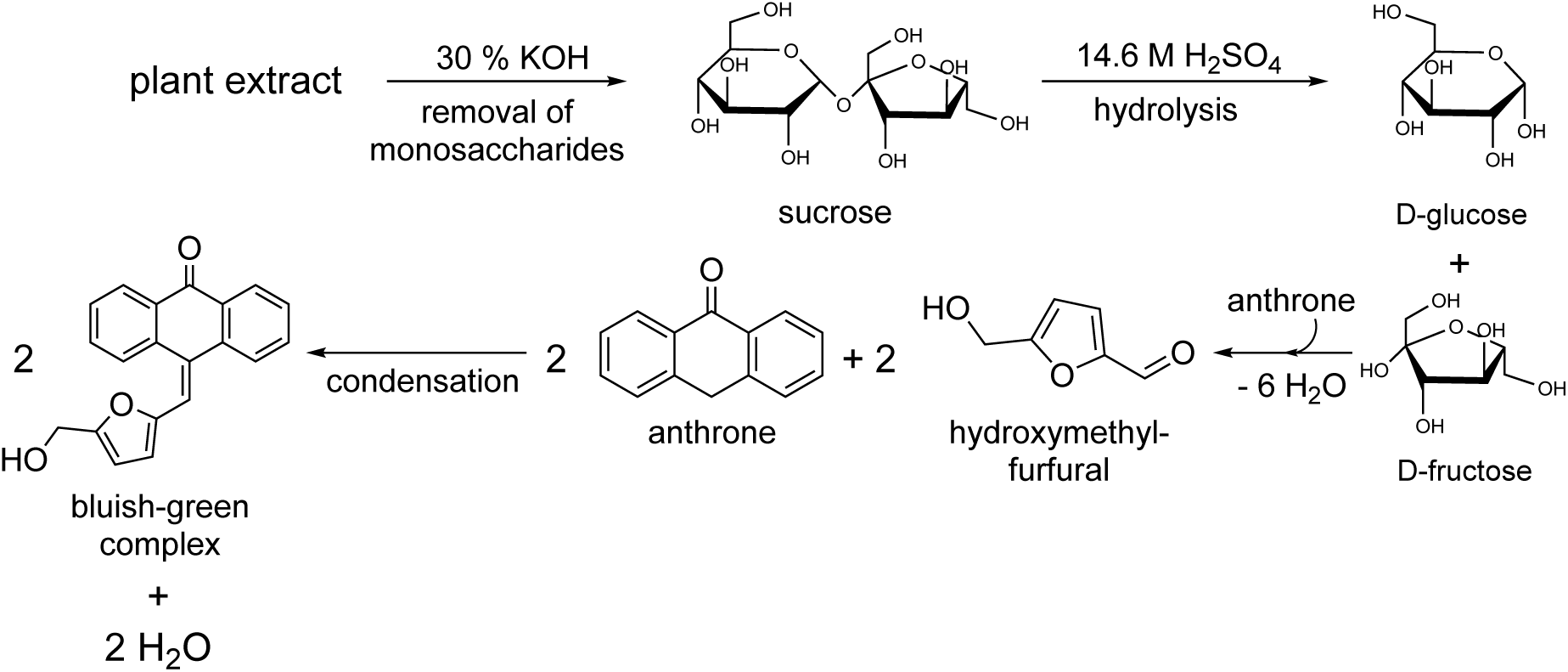
Reaction scheme for sucrose quantification via anthrone reagent. Abbreviations: H_2_SO_4_:= sulfuric acid; KOH:= potassium hydroxide. (Revvity Signal Software, 2026)

For the quantification of **glucose** and **fructose** a coupled enzyme assay was performed. Therefore plant extract was mixed with reaction buffer (100 mM imidazole-HCl (pH = 6.9), 2 mM MgCl_2_, 2 mM ATP, 1 mM NADP^+^ and 0.7 U/ml G6PDH) in a 96-well-plate and incubated until the absorbance (340 nm) remained constant (Schmidt, 1961). To determine the amount of glucose and fructose, first 1 µl hexokinase (0.5 U/µl) and then 1 µl PGI (2.5 U/µl) were added respectively. For all sugars, standards were determined in parallel and absolute amounts in samples were determined (Suppl. Fig. S3).

**Figure S3:**
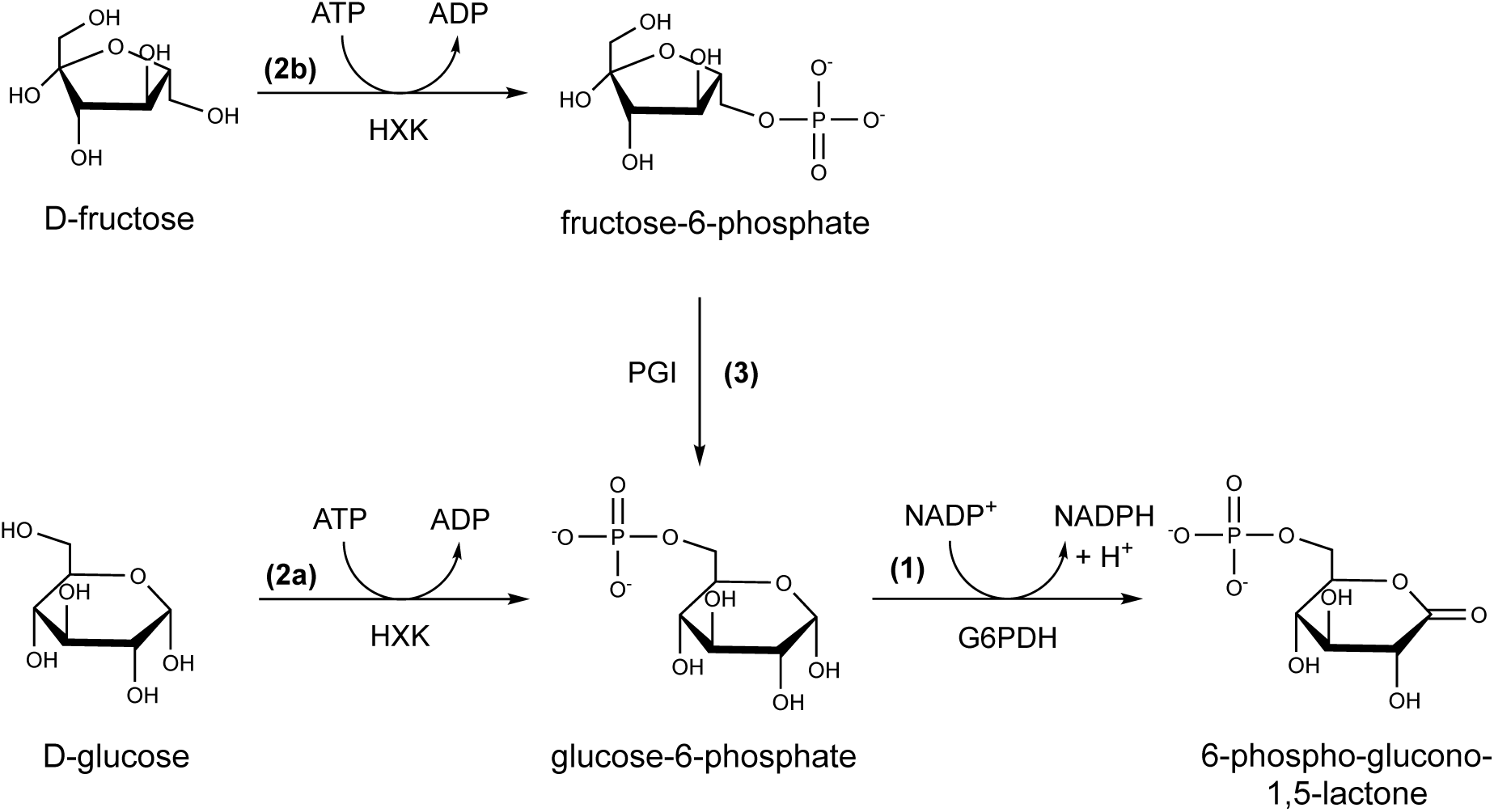
Coupled enzyme assay to quantify glucose and fructose via NADPH. G6PDH is added for background measurement (1). HXK is added to phosphorylate glucose to glucose-6-phosphate and fructose to fructose-6-phosphate for glucose quantification ((2a) and (2b)). Addition of PGI to transform fructose-6-phosphate to glucose-6-phosphate for fructose quantification (3). Abbreviations: ADP:= adenosine diphosphate; ATP:= adenosine triphosphate; G6PDH:= glucose-6-phosphate dehydrogenase; HXK:= hexokinase; NADP^+^/NADPH:= nicotinamide adenine dinucleotide phosphate; PGI:= phosphoglucoisomerase. (Revvity Signal Software, 2026)

##### Starch Quantification

For starch quantification the method previously described by Chow and Landhausser (2004) was used with following alterations. The remaining pellet of the ethanolic extraction (see A.2.1) was hydrolyzed with 0.5 M sodium hydroxide solution (NaOH) at 95 *^◦^*C for 1 h. For neutralization, 1 M acetic acid (CH_3_COOH) was added. Subsequently, the suspension was digested with 1 mg amyloglucosidase in 1 ml acetate buffer (200 mM acetic acid and 100 mM NaOH), at 55 *^◦^*C for > 14 h. The quantity of glucose obtained through the enzymatic digestion was measured through reaction of glucose and oxygen with glucose oxidase reagent (4.74 U/ml glucose oxidase, 1.7 U/ml peroxidase and 0.01% o-dianisidine-HCl in 0.5 M TRIS-HCl (pH = 7.0) and 40% (v/v) glycerin). Thereby, a pinkish complex is formed which was photometrically measured at 540 nm (Suppl. Fig. S4). Additionally, a measurement at 900 nm was conducted to assess the background, composed of residues from the plant material, if needed plant digestions were diluted.

##### G6P and F6P Quantification

**glucose-6-phosphate** and **fructose-6-phosphate** were extracted using a trichloroacetic acid (TCA)-protocol described by Hajirezaei et al. (2003) and Jelitto et al. (1992) with the following adaptations. Ground plant material (fresh mass, FM), was incubated with 16% (w/v) TCA in diethyl ether (DE) at −80 *^◦^*C for 15 min. In the next step, 16% (w/v) TCA and 5 mM ethylene glycol-bis(β-aminoethyl ether)-N,N,N’,N’-tetraacetic acid (EGTA) (in water), were added and samples incubated for 3 h on ice while occasionally shaken. After centrifugation the ether phase was removed and each sample washed three times with DE. The washed aqueous solution was neutralized by adding 1 M triethanolamin in 5 M KOH. Then a cycling assay was performed as described by Gibon et al. (2002), based on the findings of Nisselbaum and Green (1969), with slight alterations. Each standard and sample was mixed and incubated at 30 *^◦^*C for 30 min with an reaction buffer containing 50 mM tricine (pH = 9), 0.328 mM NADP^+^ and 11.2 U/ml G6PDH for the quantification of G6P and additionally 12.5 U/ml PGI for the F6P quantification. Afterwards, 0.5 M NaOH was added and the resulting mixture incubated at 95 *^◦^*C for 10 min to degrade remaining NADP^+^. The resulting mixture was neutralized by addition of 0.5 M HCl. Subsequently, the sample solution was mixed in a microplate with the second reaction buffer made of 294 mM tricine, 14.7 mM MgCl_2_, 14.7 mM EDTA, 34.9 U/ml G6PDH and 8.8 mM G6P, then with 6.25 mM thiazolyl blue and lastly with 25 mM phenazin-methosulfate. To determine the G6P and F6P concentrations kinetic measurements of this cycling assay were performed through photometrical measurements at 570 nm every 10 sec for 2 min (Suppl. Fig. S5). A standard curve was determined in parallel.

**Figure S4:**
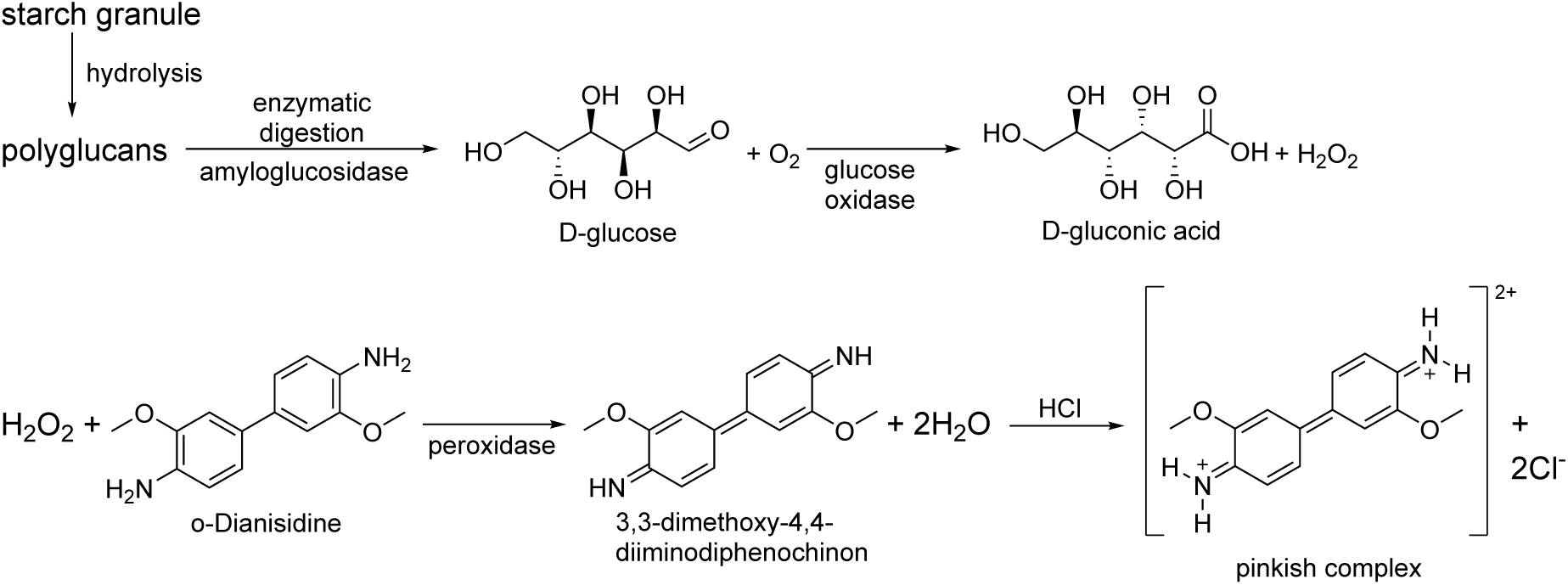
Starch degradation and glucose quantification via glucoseoxidase reagent. Abbreviations: Cl^-^:= chloride ion; H_2_O:= water; H_2_O_2_:= hydrogen peroxide; HCl:= hydrogen chloride; O_2_:= oxygen. (Revvity Signal Software, 2026)

#### A.2.2 Quantification of Enzyme Activities (*v* _max_)

##### SPS Maximum Activity

The assay to determine the maximum SPS activity was based on the assay described by J. L. A. Huber et al. (1989). Initially, ∼3 mg of dried leaf material underwent 15 min incubation on ice with extraction buffer. The extraction buffer was composed of 50 mM 4-(2-hydroxyethyl)-1-piperazine-ethanesulfonic acid potassium hydroxide (pH = 7.5) (HEPES-KOH), 10 mM MgCl_2_, 1 mM ethylenediaminetetraacetic acid (EDTA), 2.5 mM dithiothreitol (DTT), 10% glycerin and 0.1% Triton X-100. Subsequently, the suspension was centrifuged for 5 min at 21,100 g and 4 *^◦^*C. The resulting supernatant was mixed and incubated with reaction buffer at 25 *^◦^*C for 20 min. The reaction buffer contained 50 mM HEPES-KOH (pH = 7.5), 15 mM MgCl_2_, 2.5 mM DTT, 35 mM UDP-glucose, 35 mM F6P and 140 mM G6P. The reaction was halted by incubation at 95 *^◦^*C for 10 min with 30% KOH. For each sample blanks were prepared by adding 30% KOH, before the reaction buffer was added, and immediately stopping the reaction by incubation at 95 *^◦^*C (Suppl. Fig. S6). After addition of anthrone reagent the suc quantification was carried out using the anthrone assay, as described above (see section A.2.1 and Suppl. Fig. S2).

##### INVs Maximum Activities

With this assay, *v* _max_ of the soluble acidic invertase and the neutral invertase were determined as previously described by Sung et al. (1989) with following alterations. Initially, an extraction buffer was prepared composed of 50 mM HEPES-KOH pH = 7.5, 5 mM MgCl_2_, 2 mM EDTA, 1 mM phenylmethanesulfonyl fluoride solution (PMSF), 1 mM DTT, 0.1% Triton X 100 and 10% glycerin. To extract acidic and neutral invertase, the extraction buffer was added to 4 mg of dried plant material and then incubated on ice. Subsequently, two aliquots each, were used as sample solution, once for the acidic invertase and once for the neutral invertase assay. Afterwards, the samples were incubated at 30 *^◦^*C with acidic reaction buffer (20 mM sodium acetate (pH = 4.7) and 100 mM suc) for 20 min or neutral reaction buffer (20 mM HEPES KOH (pH = 7.5) and 100 mM suc) for 30 min. As neutralization buffer 1 M sodium dihydrogen phosphate (NaH_2_PO_4_) was added and the reaction stopped by immediate incubation at 95 *^◦^*C. Blanks were prepared for each sample in both assays (Suppl. Fig. S7). The amount of glc was then determined as described for the starch quantification assay (see section A.2.1 and Suppl. Fig. S4).

**Figure S5:**
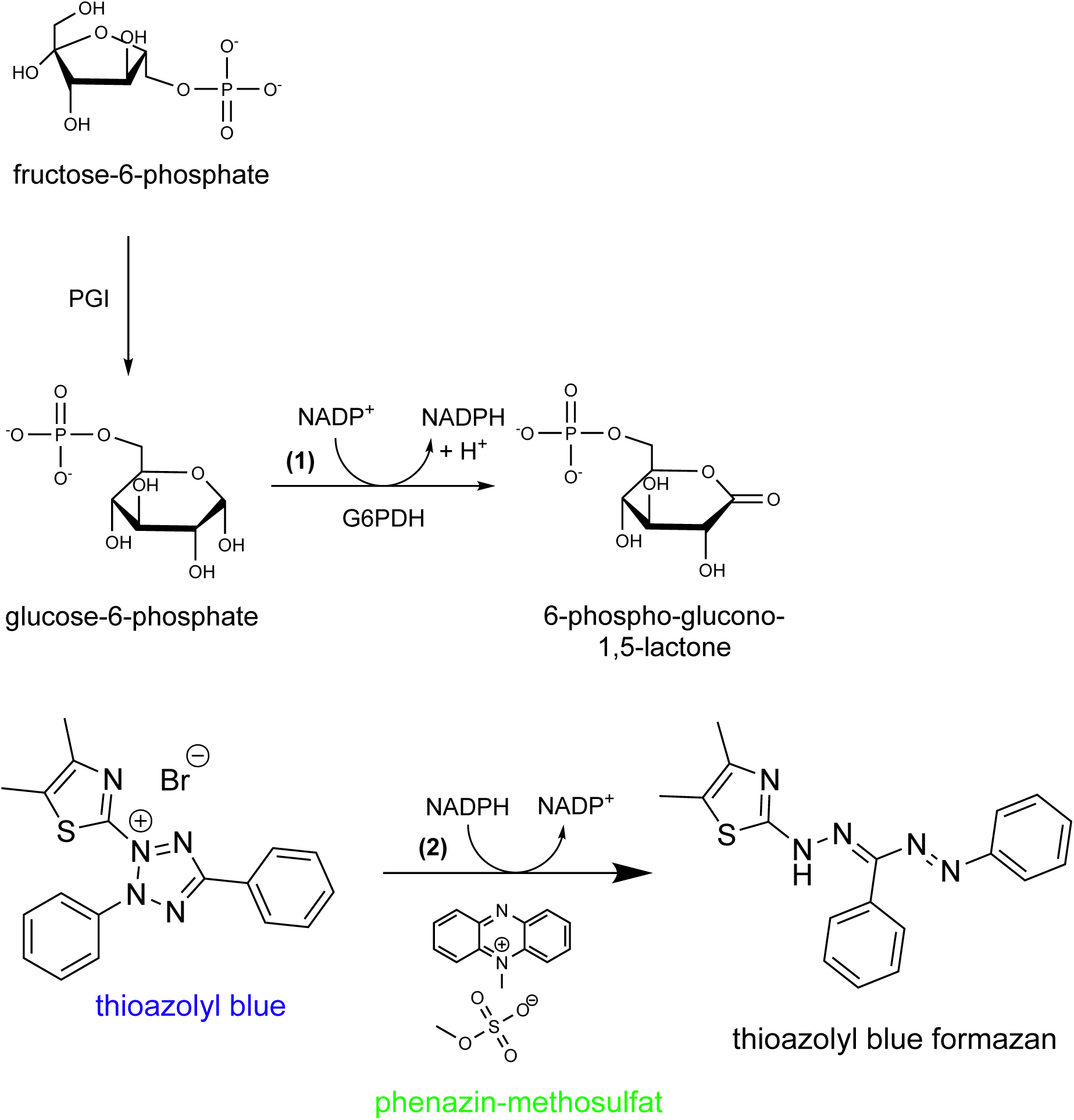
Enzyme cycling assay to quantify G6P and F6P. (1) Conversion of either G6P or F6P (through addition of PGI) conducted by G6PDH to 6-phospho-glucono-1,5-lactone generating NADPH as side product. (2) Starting the cycling assay by addition of thioazolyl blue, phenazin-methosulfat and G6PDH. Abbreviations: Br^-^:= bromide ion; G6PDH:= glucose-6-phosphate dehydrogenase; NADP^+^/NADPH:= nicotinamide adenine dinucleotide phosphate; PGI:= phosphoglucoisomerase. (Revvity Signal Software, 2026)

**Figure S6:**
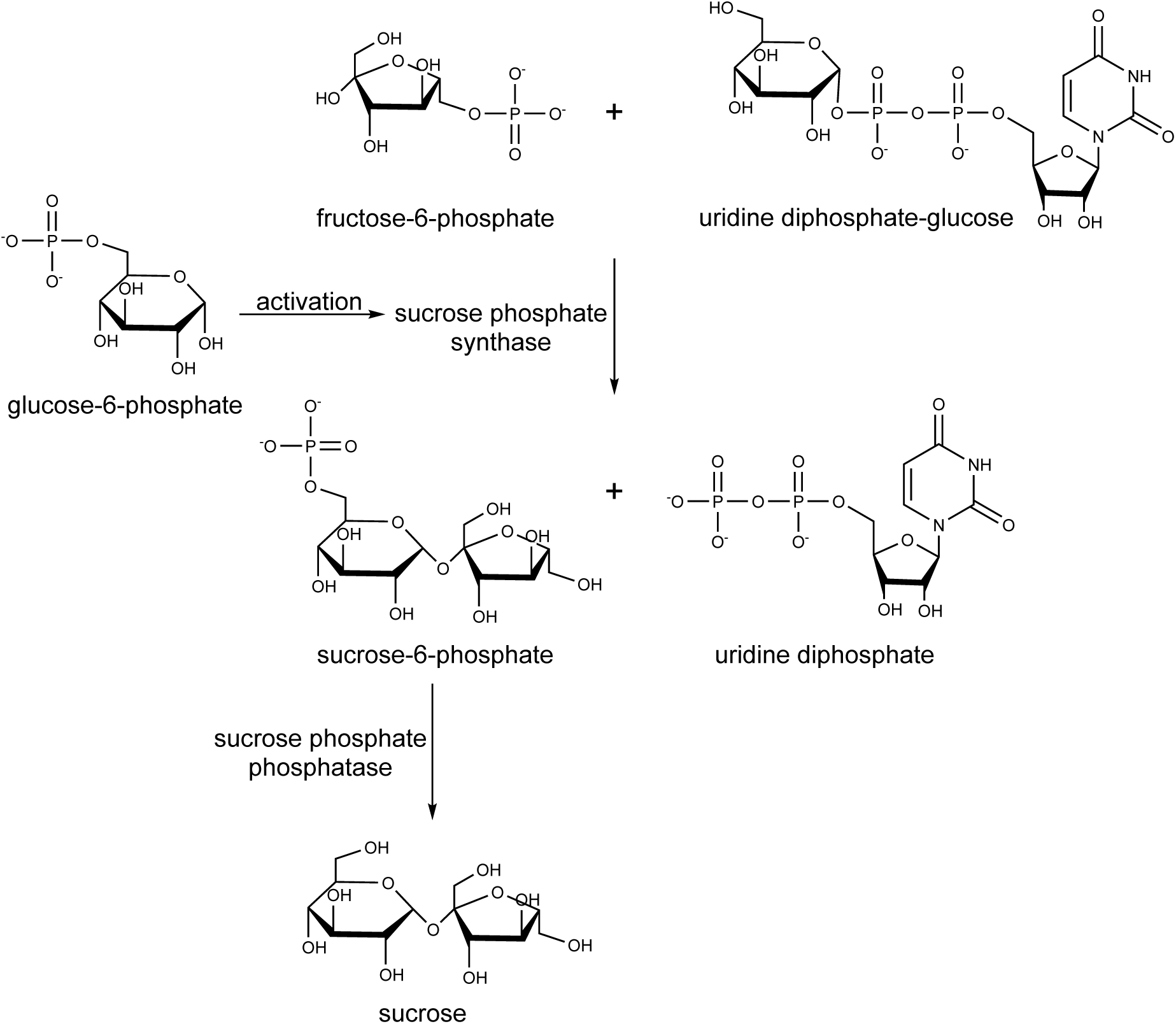
Formation of sucrose via SPS and SPP. Abbreviations: SPP:= sucrose-phosphate phosphatase; SPS: sucrose-phosphate synthase. (Revvity Signal Software, 2026)

**Figure S7:**
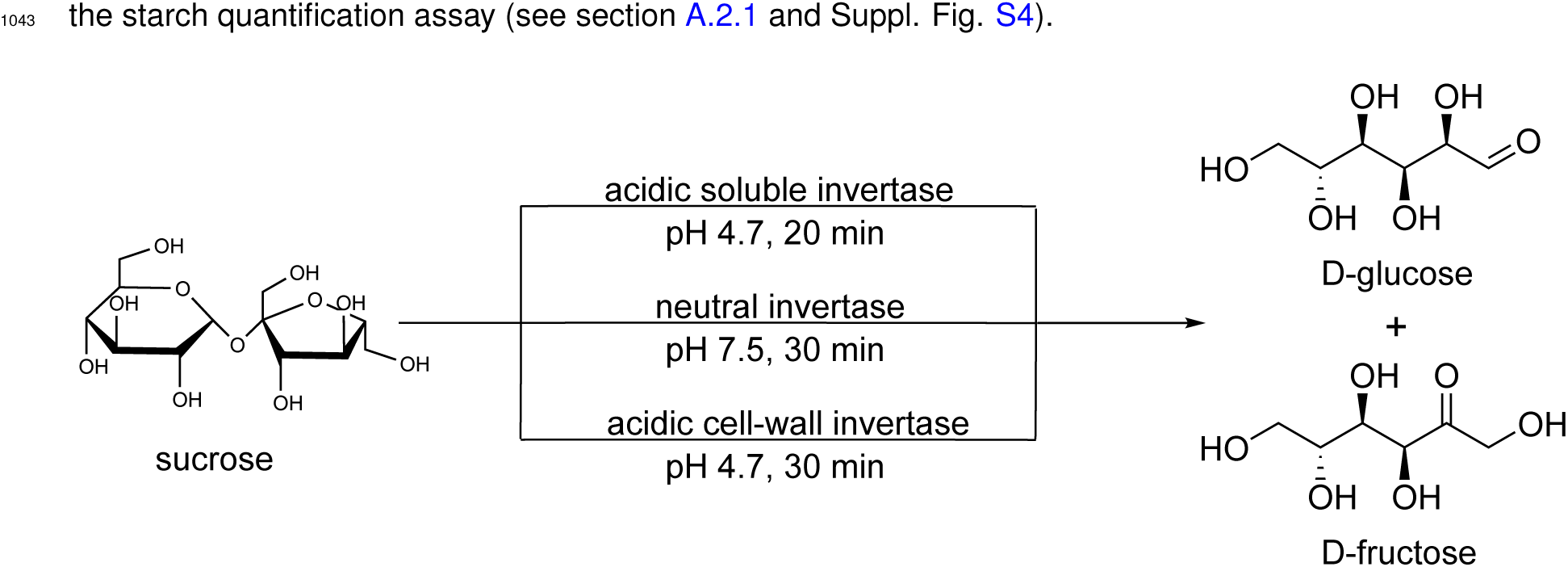
Cleavage of sucrose via INVs to glucose and fructose. Determination of acidic and neutral invertase activity. For cell-wall invertase only one replicate was measured (therefore it is not included in the presented data). Abbreviations: INV:= invertase. (Revvity Signal Software, 2026)

##### HXK Maximum Activities

The maximum hexokinases activities were determined with the assay described by Wiese et al. (1999) with slight alterations. First, an extraction buffer was prepared containing 50 mM TRIS-HCl (pH = 8.0), 0.5 mM MgCl_2_, 1 mM EDTA, 0.1% Triton X-100 and 2.5 mM DTT. Next, 2.5 mg of dried plant material were incubated with the extraction buffer for 15 min on ice. The reaction buffer was prepared of a stock solution containing 100 mM HEPES-KOH (pH = 7.5), 10 mM MgCl_2_, 100 mM NADP^+^ and G6PDH (0.5 U/µl) for the glucokinase measurement, with the addition of PGI (0.5 U/µl) for the fructokinase measurements. For sample measurements 100 mM ATP was added, while for the blanks H_2_O was added instead. In a microplate reaction buffer was mixed with sample extract and 50 mM glc or frc respectively. The concentration of NADPH, which is obtained as side product, was photometrically measured at 340 nm every minute over 15 min. The maximum activities *v* _max_ were then determined based on the resulting slopes of the kinetic measurements and the law of Lambert-Beer (Wypych, 2015). The same reaction cascade as described in section A.2.1 was used to determine GLCK and FRCK activity by supplying glc and frc as substrates, ATP as activator and G6PDH to measure the kinetics of NADPH formation (Suppl. Fig. S3).

#### A.2.3 Augmented Neural ODE (ANODE) Motivation

The system under study comprises six measured metabolite pools: fructose-6-phosphate (F6P), glucose-6-phosphate (G6P), glucose (glc), sucrose (suc), fructose (frc), and starch (Sta). A standard neural ODE (NODE) operating solely on these observed variables assumes the dynamics are Markovian in the measurement space, i.e. that the observed concentrations fully determine the system’s future trajectory. Initial experiments showed that this assumption does not hold, motivating the use of an *augmented* NODE (ANODE) (Dupont et al., 2019) in which the observed state is lifted into a higher-dimensional latent space containing *n*_hid_ = 1 additional hidden (latent) variables. The ODE is then solved in this augmented space, and the solution is projected back onto the observable dimensions for comparison with data.

#### A.2.4 Model architecture

Let **y**_obs_(*t*) *∈* R*^n^*^obs^, *n*_obs_ = 6, denote the vector of observed metabolite concentrations at time *t*. The augmented state is

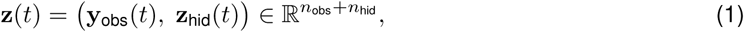

where **z**_hid_(*t*) *∈* R*^n^*^hid^ represents latent biochemical context not directly observed.

##### Encoder and initial conditions

The initial latent state is constructed by concatenating the observed initial concentrations **y**_obs_(*t*_0_) with a learned parameter vector **z**_0_ *∈* R*^n^*^hid^:

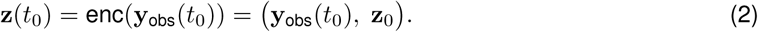

The parameters **z**_0_ are optimized jointly with the rest of the model.

##### Dynamics

The right-hand side of the ODE is decomposed into a *biology-informed* part, governed by a known stoichiometric structure, and a purely data-driven *Markovian correction* for the hidden variables:

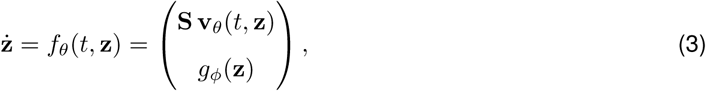

where **S** *∈* R*^n^*^obs^ *^×^*^12^ is the stoichiometric matrix of the core carbon metabolism (fixed by prior knowledge), **v***_θ_* is the flux vector (parameterized by an multi-layer perceptron (MLP)), and *g_ϕ_*is the hidden-state dynamics MLP.

##### Stoichiometric structure

The stoichiometric matrix encodes the following equations:

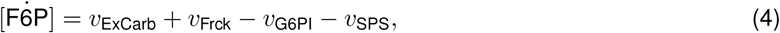

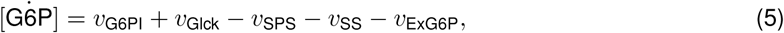

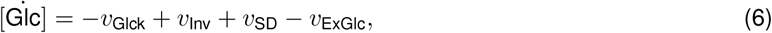

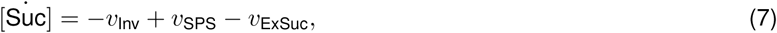

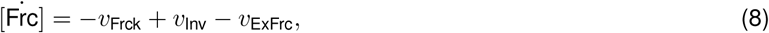

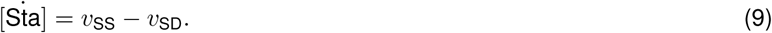

The twelve fluxes are: sucrose phosphate synthase (SPS), invertase (INV), glucokinase (GLCK), fructokinase (FRCK), glucose-6-phosphate isomerase (GPI), starch synthase (SS), starch degradation (SD), and five exchange fluxes (ExCarb, ExGlc, ExSuc, ExG6P, ExFrc), of which only ExCarb and ExGlc are assumed to be reversible.

##### Flux MLP

The flux vector **v***_θ_*is computed by an MLP

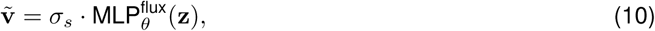

with input dimension *n*_obs_ + *n*_hid_ = 7, output dimension 12, hidden width 64, depth 6, and Softplus activations. A global output scale *σ_s_*, initially 0.1, was introduced to prevent initial flux magnitudes from driving the system into stiff regimes. Enzyme-capacity constraints are incorporated by multiplying selected fluxes by time-varying maximal rates *V* ^max^(*t*) estimated from separate enzyme-activity measurements via cubic splines, applying a saturating normalization 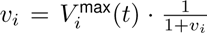. Fluxes constrained to be non-negative (starch synthesis/degradation, export fluxes) are passed through an absolute-value operation.

##### Hidden-state MLP

The dynamics of the hidden variables are governed by a second MLP, *g_ϕ_*(**z**), with input dimension 7, output dimension 1, hidden width 64, depth 6, and Softplus activations.

##### Decoder and soft-to-hard annealing

For loss computation, the latent trajectory must be projected back to the observable space. A *hard* decoder simply reads off the first *n*_obs_ components of **z**, which is the physically correct projection. However, early in training the hidden-variable gradients vanish under the hard decoder because changes to **z**_hid_ have no direct effect on the observable loss — the hidden dimensions are cut off before the comparison with data. This gradient starvation prevents the hidden state from learning a useful representation during the initial phase of training.

To circumvent this, we introduce a *soft* decoder, a linear layer *D_ψ_*: R*^n^*^obs+^*^n^*^hid^ *→* R*^n^*^obs^, which mixes all latent dimensions and thus propagates gradients into **z**_hid_. The effective decoder is a convex combination controlled by an annealing coefficient *α ∈* [0, 1]:

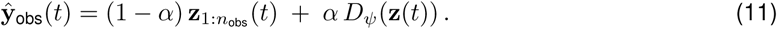

At the start of training *α ≈* 1 (soft decoder dominates, hidden gradients flow freely); as training progresses *α* is annealed towards zero so that by post-training only the physically meaningful hard projection is used. The schedule follows a sigmoid:

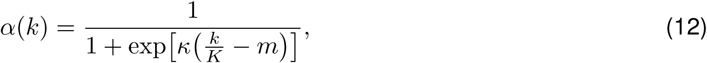

where *k* is the global training step, *K* is the total number of pre-training steps, *κ* = 20 controls the steepness of the transition, and *m* = 0.4 sets the midpoint.

#### A.2.5 ODE integration

The ANODE is implemented using equinox integrated with the diffrax library (Kidger & Garcia, 2021). During pre-training, the implicit Kvaerno5 solver (a stiffly-stable Runge–Kutta method) is used to handle the transient stiffness that arises from early, poorly-constrained MLP parameterizations. Once pre-training is complete and the dynamics have stabilized, the explicit Tsit5 solver (an efficient 5th-order Runge–Kutta method) is used throughout post-training for faster computation. In both phases, step sizes are controlled by a PID controller with relative and absolute tolerances of 10*^−^*^4^ and 10*^−^*^6^, respectively, and a maximum of 8,192 solver steps per integration.

#### A.2.6 Training procedure

##### Loss

The model is fitted by minimizing the mean-squared error between predicted and measured metabolite trajectories, normalized by the per-metabolite standard deviation estimated from the data:

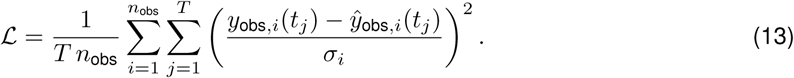

##### Curriculum pre-training

Training begins with a curriculum that starts on a short prefix of the time series and progressively extends to the full integration window (fractions 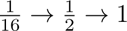 with 500, 500, and 20,000 gradient steps, respectively). This prevents the model from being overwhelmed by long-horizon stiff dynamics early in training. The AdaBelief optimizer (Zhuang et al., 2020) with learning rate 10*^−^*^4^ is used throughout pre-training.

##### Post-training

Following pre-training, *α* is fixed to zero (hard decoder only) and the model is further optimized over the full time series for up to 200,000 steps, again with a brief curriculum warm-up (500 and 500 steps at fractions 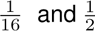). The AdamW optimizer (Loshchilov & Hutter, 2017) with learning rate 10*^−^*^4^ and reduced momentum *β*_1_ = 0.5 is used; the lower momentum prevents the optimizer from overshooting into stiff parameter regions, which is particularly harmful for NODE training (Kidger, 2021).

##### Gradient handling

Gradients are clipped to a global *ℓ*_2_ norm of 1.0 at every step. When the ODE solver fails (e.g. due to a stiff parameterization exceeding the maximum step count), the resulting NaN gradients are replaced with zeros before the optimizer update, preventing corruption of the optimizer state.

##### Early stopping

Post-training terminates early once the loss falls below a threshold of 10*^−^*^4^. The model checkpoint with the lowest post-training loss is retained.

#### A.2.7 Ensemble training

To account for sensitivity to random initialization inherent in neural ODE training, we trained an ensemble of 100 independently initialized models for each of the three conditions: ambient temperature control, one day of chilling night adaptation, and seven days of chilling night acclimation. Models with a loss above 1 *×* 10*^−^*^1^ were deemed ill-fitted and filtered out of the analysis. Each model is initialized with a distinct random seed; the best-performing checkpoint (minimum post-training loss) from each seed is retained. To account for the fact that the control and chilling night day 1 conditions started at the same initial conditions, we did a post-training step where we trained chilling night day 1 to match the fluxes of control at time point 00 h. Ensemble predictions of fluxes and concentrations are reported as means *±* SEM across the models.

### A.3 Supplemental Results

#### A.3.1 Photosynthesis

**Figure S8:**
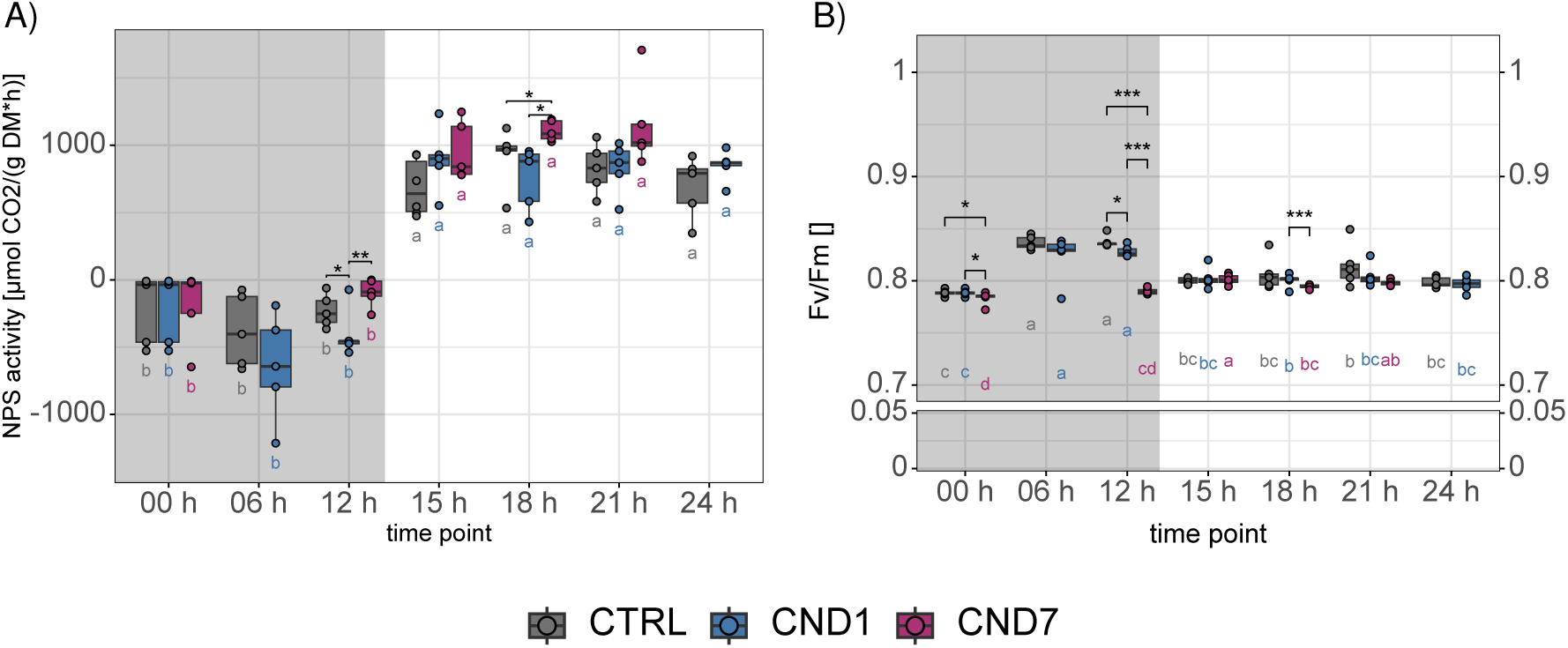
Photosynthetic activity and Fv/Fm (maximum potential quantum efficiency of Photosystem II) during chilling night events. **A)** The photosynthetic activity (NPS) and **B)** the photosynthetic capacity of photosystem II (F_v_/F_m_) in i) control (CTRL, gray, 18 *^◦^*C night and 22 *^◦^*C day), ii) chilling night day one (CND1, blue, 4 *^◦^*C night and 22 *^◦^*C day) and iii) chilling night day seven (CND7, magenta, 4 *^◦^*C nights and 22 *^◦^*C days). The photosynthetic activity (NPS) in µmol CO_2_ per gram DM per hour and (F_v_/F_m_) are plotted against the TPs [h]. The dark gray area indicates the nighttime, and light gray area the daytime. Five replicates were measured for each condition at each TP except for day seven 06 h and 24 h TP (no harvest). A t-test was performed between conditions at each TP (* p *<* 0.05, ** p *<* 0.01, *** p *<* 0.001). A one-way ANOVA with a Tukey HSD as a post hoc test (p *<* 0.05) was performed within each condition across TPs and is depicted as a CLD below each boxplot in the corresponding condition color. Abbreviations: ANOVA:= analysis of variance; CLD:= compact letter display; CND1:= chilling night day 1; CND7:= chilling night day 7; CTRL:= control; Fv/Fm:= photosynthetic capacity of Photosystem II; HSD:= honestly significant difference; NPS:= net photosynthesis; TP= time point. R-package:Shuangbin Xu et al. (2021)

#### A.3.2 Sugar Phosphates

**Figure S9:**
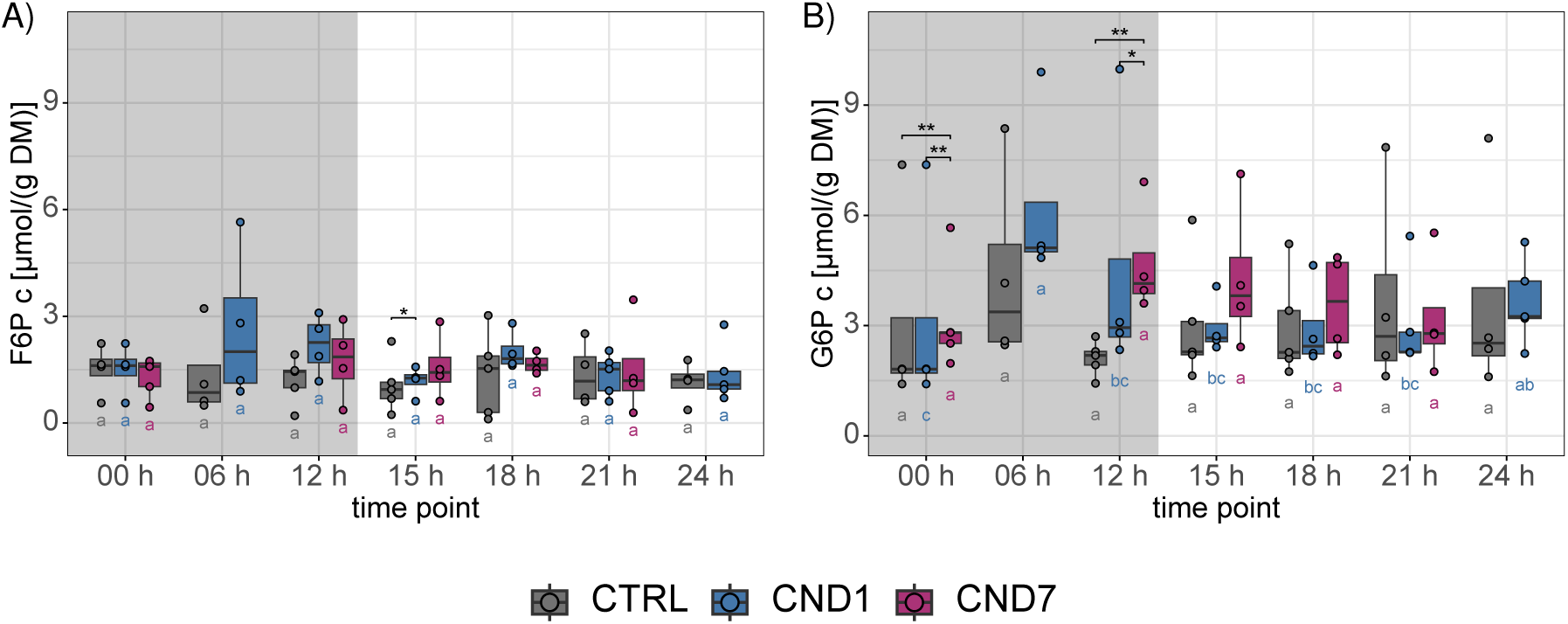
Sugar phosphate concentrations during chilling night events. **A)** Glucose-6-phosphate (G6P) and **B)** fructose-6-phosphate (F6P) concentrations in i) control (CTRL, gray, 18 *^◦^*C night and 22 *^◦^*C day), ii) chilling night day one (CND1, blue, 4 *^◦^*C night and 22 *^◦^*C day) and iii) chilling night day seven (CND7, magenta, 4 *^◦^*C nights and 22 *^◦^*C days). The concentrations in µmol per gram DM are plotted against the TPs [h]. The dark gray area indicates the nighttime, and light gray area the daytime. Five replicates were measured for each condition at each TP except for day seven 06 h and 24 h TP (no harvest). A t-test was performed between conditions at each TP (* p *<* 0.05, ** p *<* 0.01, *** p *<* 0.001). A one-way ANOVA with a Tukey HSD as a post hoc test (p *<* 0.05) was performed within each condition across TPs and is depicted as a CLD below each boxplot in the corresponding condition color. Abbreviations: ANOVA:= analysis of variance; c:=concentration; CLD:= compact letter display; CND1:= chilling night day 1; CND7:= chilling night day 7; CTRL:= control; F6P:= fructose-6-phosphate; G6P:= glucose-6-phosphate; HSD=: honestly significant difference; TP:= time point.

#### A.3.3 INV and HXK maximum enzyme activity *v* _max_

**Figure S10:**
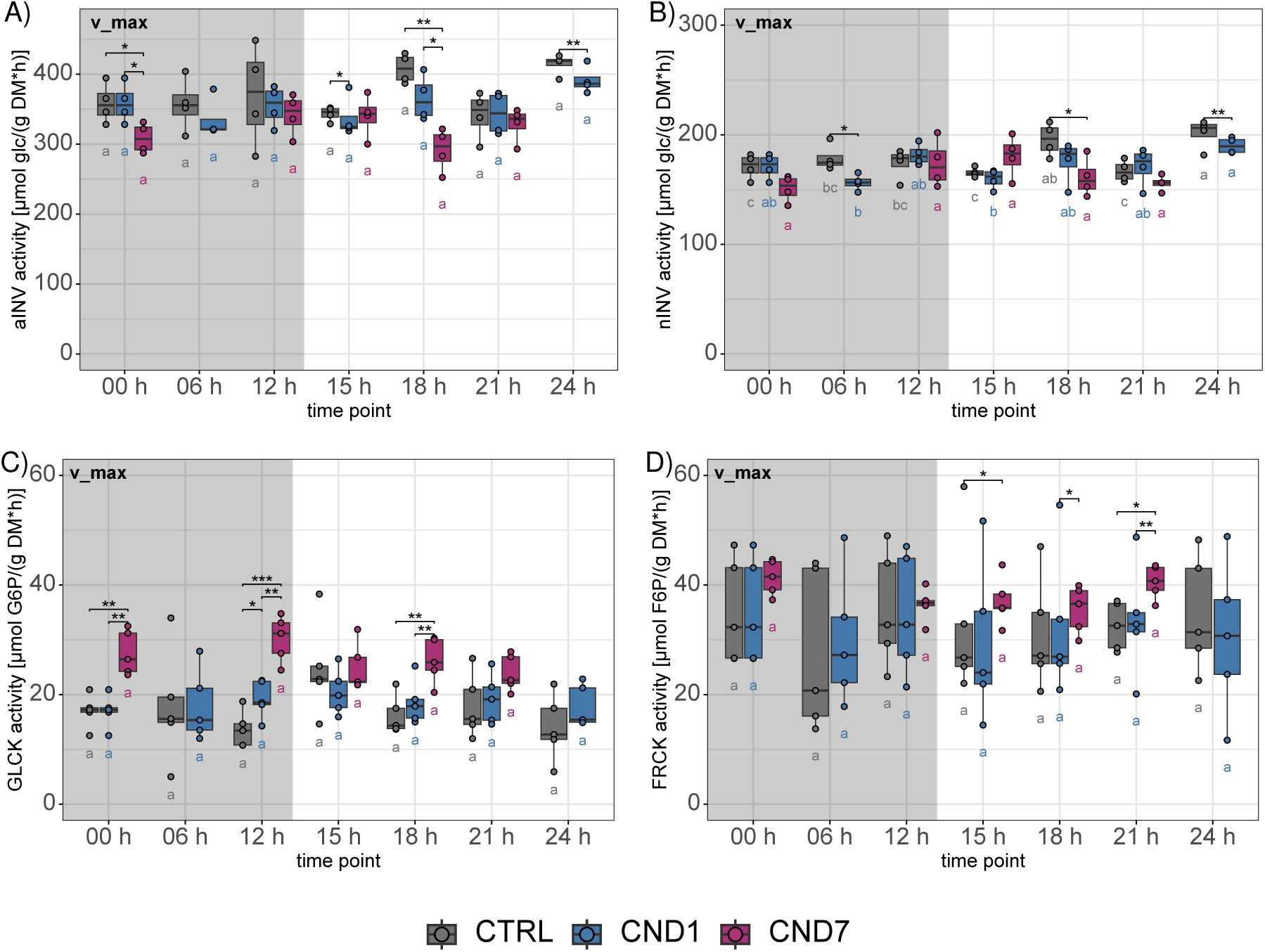
Maximum invertase and kinase activity (*v* _max_) during chilling night events. Maximum activity of **A)** acidic invertase (aINV), **B)** neutral invertase (nINV), **C)** glucokinase (GLCK) and **D)** fructokinase (FRCK) in i) control (CTRL, gray, 18 *^◦^*C night and 22 *^◦^*C day), ii) chilling night day one (CND1, blue, 4 *^◦^*C night and 22 *^◦^*C day) and iii) chilling night day seven (CND7, magenta, 4 *^◦^*C nights and 22 *^◦^*C days). The *v* _max_ in µmol per gram DM per hour are plotted against the TPs [h]. The dark gray area indicates the nighttime, and light gray area the daytime. Five replicates were measured for each condition at each TP except for day seven 06 h and 24 h TP (no harvest). A t-test was performed between conditions at each TP (* p *<* 0.05, ** p *<* 0.01, *** p *<* 0.001). A one-way ANOVA with a Tukey HSD as a post hoc test (p *<* 0.05) was performed within each condition across TPs and is depicted as a CLD below each boxplot in the corresponding condition color. Abbreviations: ANOVA:= analysis of variance; CLD:= compact letter display; CND1:= chilling night day 1; CND7:= chilling night day 7; CTRL:= control; F6P:= fructose-6-phosphate; FRCK:= fructokinase; G6P:= glucose-6-phosphate; glc:= glucose; GLCK:= glucokinase; HSD=: honestly significant difference; n/aINV:= neutral/acidic invertase; TP:= time point.

#### A.3.4 Arrhenius corrected maximum enzyme activity (*v* _max_) of INV and HXK

**Figure S11:**
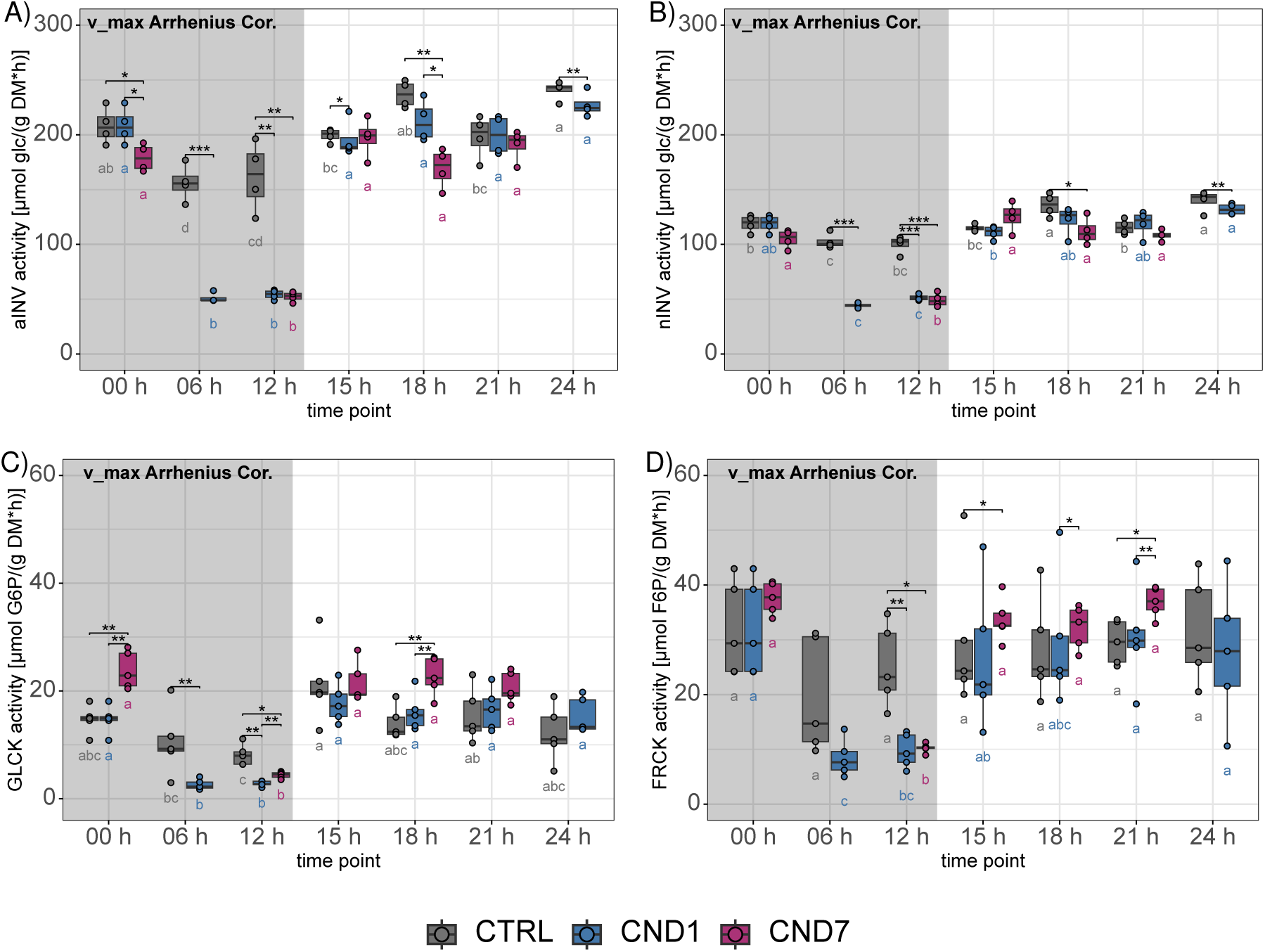
Arrhenius corrected maximum invertase and kinase activity (*v* _max_) during chilling night events. By Arrhenius correction (Arrhenius, 1889a, 1889b; Laidler, 1984) temperature adapted maximum activity of **A)** acidic invertase (aINV), **B)** neutral invertase (nINV), **C)** glucokinase (GLCK) and **D)** fructokinase (FRCK) in i) control (CTRL, gray, 18 *^◦^*C night and 22 *^◦^*C day), ii) chilling night day one (CND1, blue, 4 *^◦^*C night and 22 *^◦^*C day) and iii) chilling night day seven (CND7, magenta, 4 *^◦^*C nights and 22 *^◦^*C days). The Arrhenius corrected *v* _max_ in µmol per gram DM per hour are plotted against the TPs [h]. The dark gray area indicates the nighttime, and light gray area the daytime. Five replicates were measured for each condition at each TP except for day seven 06 h and 24 h TP (no harvest). A t-test was performed between conditions at each TP (* p *<* 0.05, ** p *<* 0.01, *** p *<* 0.001). A one-way ANOVA with a Tukey HSD as a post hoc test (p *<* 0.05) was performed within each condition across TPs and is depicted as a CLD below each boxplot in the corresponding condition color. Abbreviations: ANOVA:= analysis of variance; CLD:= compact letter display; CND1:= chilling night day 1; CND7:= chilling night day 7; CTRL:= control; F6P:= fructose-6-phosphate; FRCK:= fructokinase; G6P:= glucose-6-phosphate; glc:= glucose; GLCK:= glucokinase; HSD=: honestly significant difference; n/aINV:= neutral acidic invertase; TP:=time point.

#### A.3.5 Fructose

**Figure S12:**
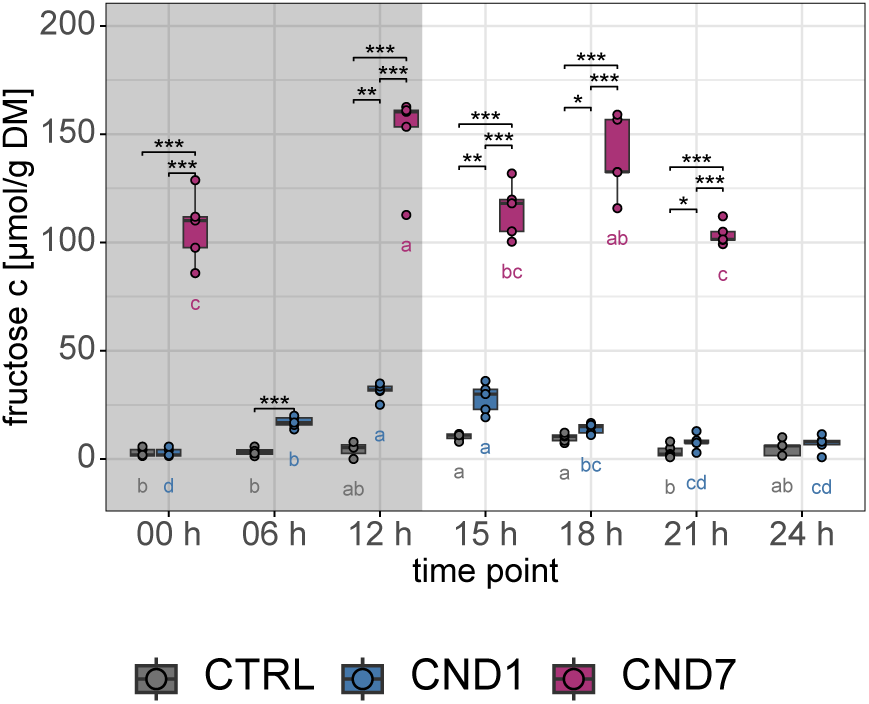
Fructose concentration in the course of chilling night events. Fructose concentrations in i) control (CTRL, gray, 18 *^◦^*C night and 22 *^◦^*C day), ii) chilling night day one (CND1, blue, 4 *^◦^*C night and 22 *^◦^*C day) and iii) chilling night day seven (CND7, magenta, 4 *^◦^*C nights and 22 *^◦^*C days). The concentrations in µmol per gram DM and are plotted against the TPs [h]. The dark gray area indicates the nighttime, and light gray area the daytime. Five replicates were measured for each condition at each TP except for day seven 06 h and 24 h TP (no harvest). A t-test was performed between conditions at each TP (* p *<* 0.05, ** p *<* 0.01, *** p *<* 0.001). A one-way ANOVA with a Tukey HSD as a post hoc test (p *<* 0.05) was performed within each condition across TPs and is depicted as a CLD below each boxplot in the corresponding condition color. Abbreviations: CLD:= compact letter display; CND1:= chilling night day 1; CND7:= chilling night day 7; CTRL:= control

#### A.3.6 Correlation Plots

**Figure S13:**
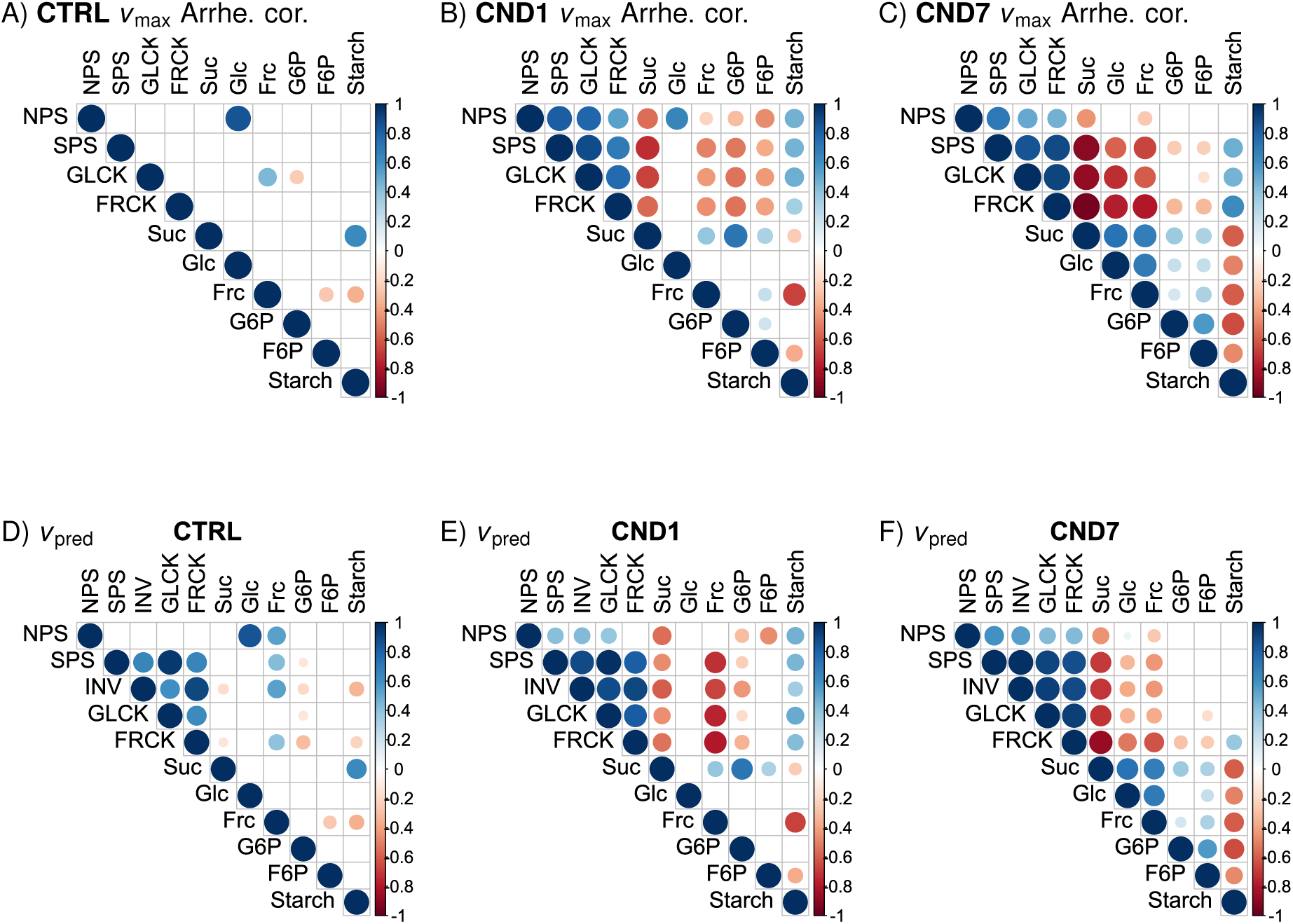
Correlation between net photosynthesis, metabolite concentrations and maximum enzyme activities (*v* _max_) or modeled enzyme rates (*v* _pred_). Correlation plots (linear correlation (Pearson)) of all metabolites and NPS for each time series with **A-C)** maximum enzyme activities (*v* _max_) and **D-F)** modeled rates *v* _pred_. Diurnal correlations are in A) and D) for control, in B) and E) for chilling night day 1, in C) and F) for chilling night day 7. Depicted are correlations with t-test p *<* 0.05, the circle size depicting the significance level with red representing negative and blue positive correlations. Abbreviations: Arrhe. cor.:= Arrhenius corrected; CND1:= chilling night day 1; CND7:= chilling night day 7; CTRL:= control; DM:= dry mass; F6P:= fructose-6-phosphate; frc:= fructose; FRCK:= fructokinase; G6P:= glucose-6-phosphate; glc:= glucose; GLCK:= glucokinase; n/aINV:= neutral/acidic invertase; NPS:= net photosynthesis; SPS:= sucrose phosphate synthase; suc:= sucrose;

#### A.3.7 Starch Metabolism

**Figure S14:**
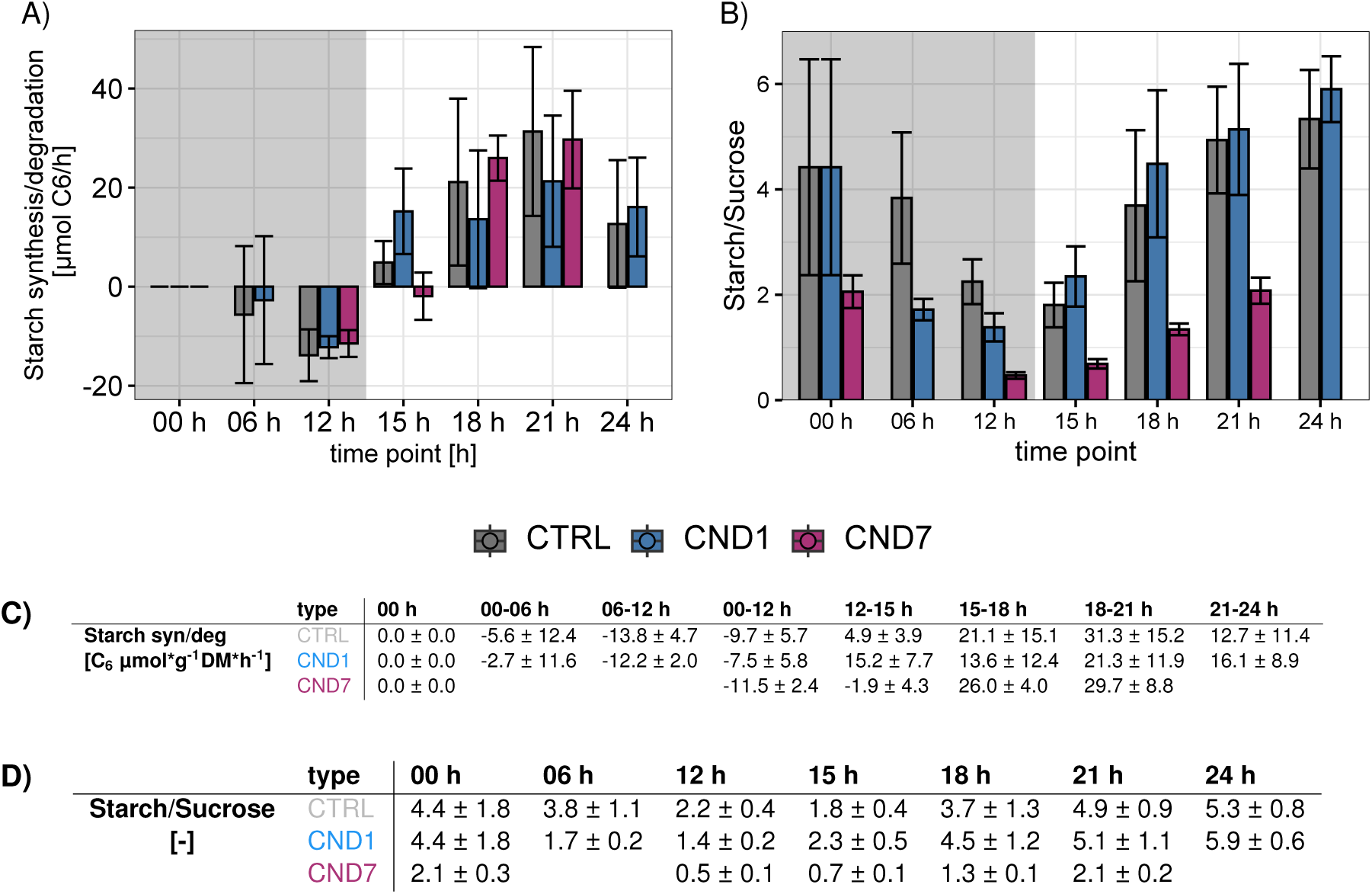
Starch synthesis/degradation and starch/suc ratio during chilling night events. **A)** Starch synthesis/degradation was calculated as concentration difference between two consecutive time points divided by the time between the time points. The law of error propagation was applied to calculate the errors for each time point. **B)** The ratio from starch to sucrose (C_6_ equivalents) was calculated from the mean concentrations per time point and condition. The law of error propagation was applied to calculate the errors for each time point and condition. **C)** All calculated values of starch synthesis/degradation *±* standard error are listed in the table (for better comparison of night degradation, difference between 00 h and 12 h were calculated for CTRL and CND1, additionally). **D)** All calculated values of starch/sucrose ratio (with sucrose calculated to C_6_ equivalents) are listed for each time point *±* standard error. Starch syn/deg and starch/sucrose ratio in i) control (CTRL, gray, 18 *^◦^*C night and 22 *^◦^*C day), ii) chilling night day one (CND1, blue, 4 *^◦^*C night and 22 *^◦^*C day) and iii) chilling night day seven (CND7, magenta, 4 *^◦^*C nights and 22 *^◦^*C days). The dark gray area indicates the nighttime, and light gray area the daytime. Abbreviations: CTRL:= control; CND1:= chilling night day 1; CND7:= chilling night day 7; syn/deg:= synthesis/degradation.

#### A.3.8 Overview Data

**Figure S15:**
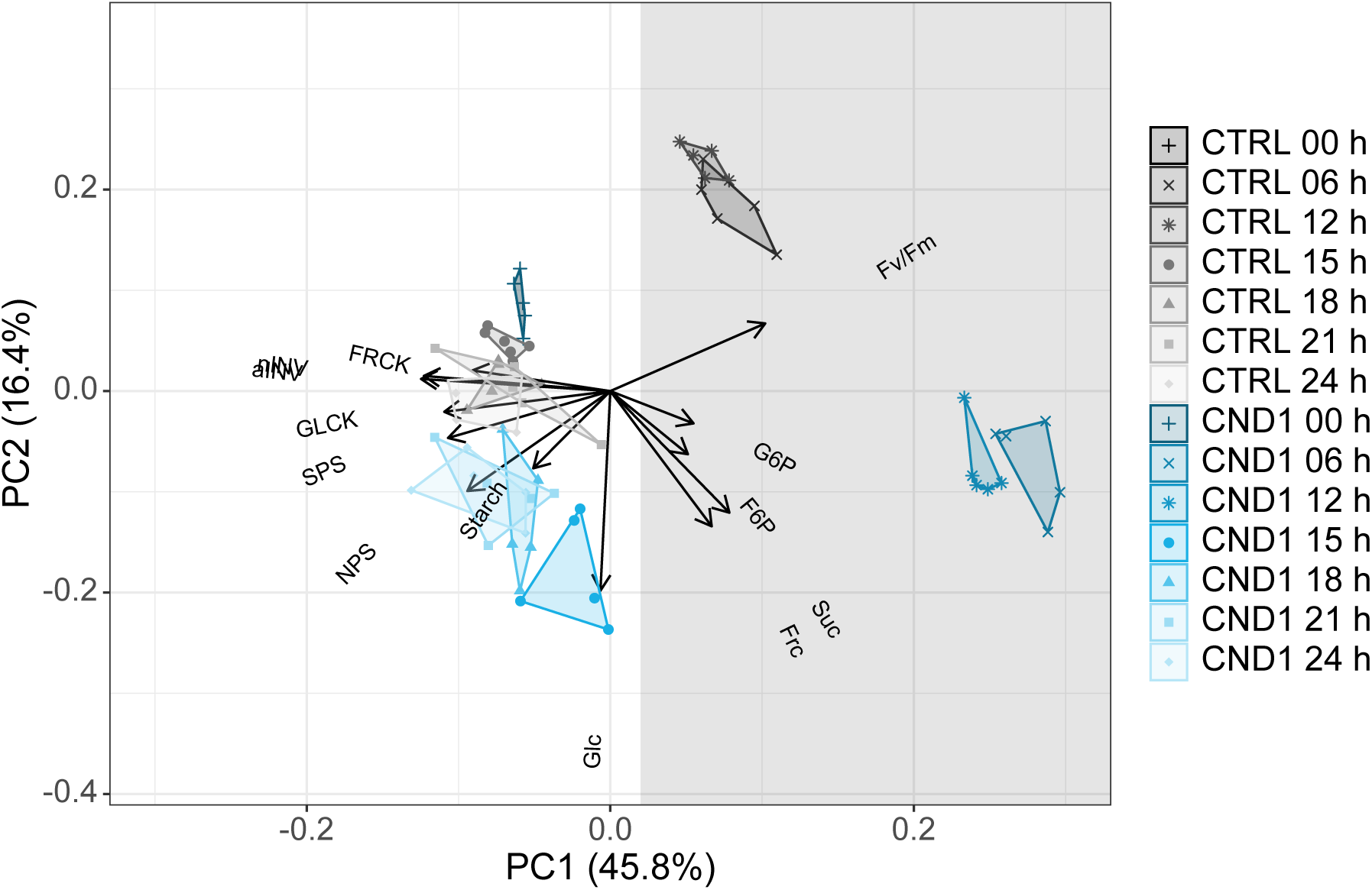
Principal component analysis for control and chilling night day one data. When plotting only the data of the control vs. chilling night day one still the day time points (15 h to 24 h) are only just separated form each other. Night time points (6 h and 12 h) are already clearly separated. Abbreviations: CND1:= chilling night day 1; CND7:= chilling night day 7; CTRL:= control; F6P:= fructose-6-phosphate; frc:= fructose; FRCK:= fructokinase; Fv/Fm:= maximum potential quantum efficiency of Photosystem II; G6P:= glucose-6-phosphate; glc:= glucose; GLCK:= glucokinase; md:=midday; n/aINV:= neutral/acidic invertase; NPS:= net photosynthesis; PC:= principal component; suc:= sucrose; SPS:= sucrose phosphate synthase.

**Table S1:**
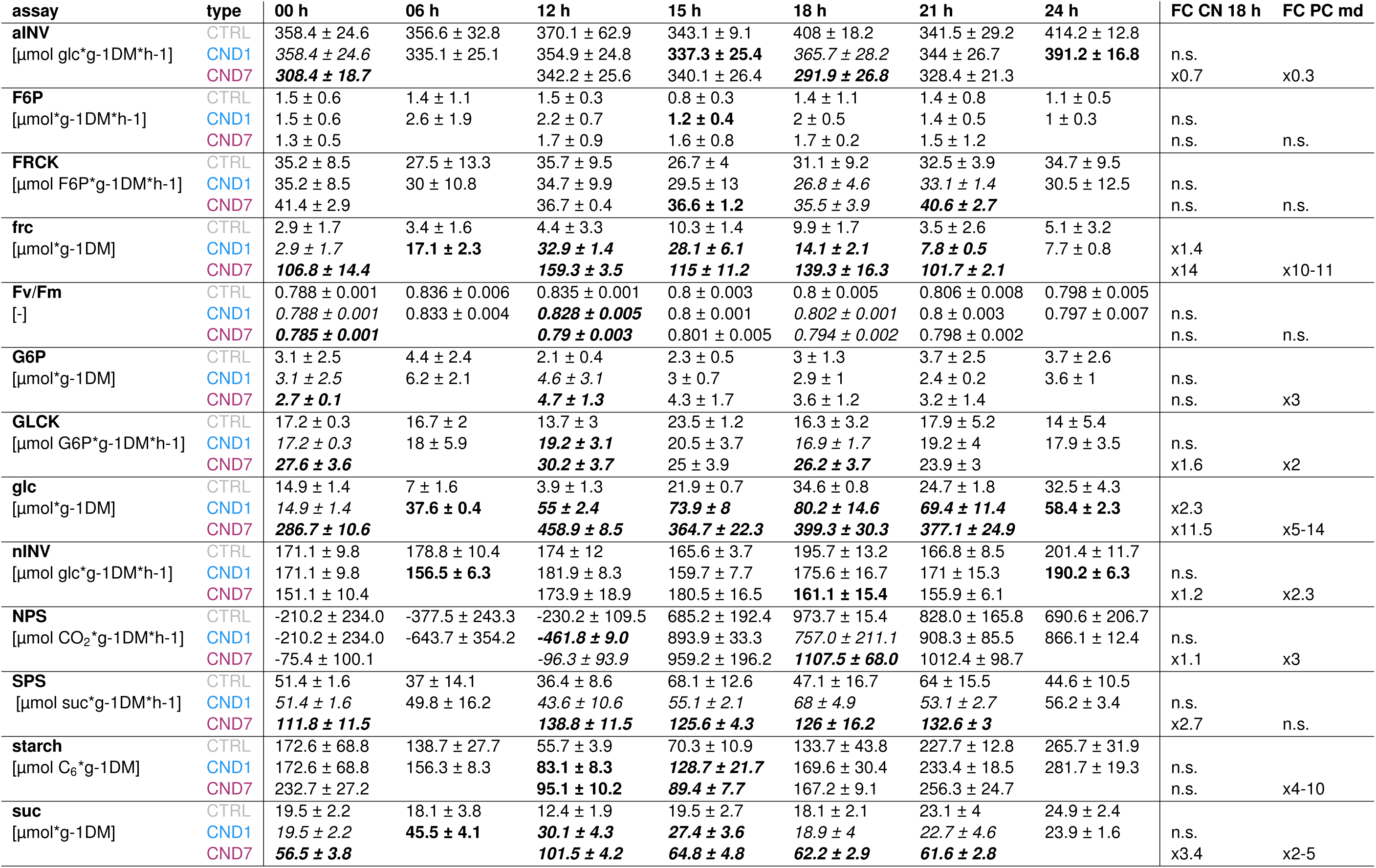
Carbohydrate concentrations and related maximum enzymes activities *v* _max_ for control and chilling night conditions. Mean values ± standard deviation for all measured carbohydrate concentrations in [µmol·g^-1^ DM] and related maximum enzyme activities (*v* _max_) in [µmol·g^-1^ DM·h^-1^
]. Significant differences from the control (CTRL) are depicted **bold**. When there is a significant difference between the chilling night day 1 (CND1) and the chilling night day 7 (CND7) they are both depicted in *italics*. FC CN 18 h describes the mean fold change from CTRL to CND7 for the 18 h (midday) time point. FC PC represents mean fold change from midday (md) day 7 of permanent cold condition (4 *^◦^*C) normalized to respective control investigated by (Kitashova et al., 2021, 2023). Abbreviations: aINV:= acidic invertase; CN:= chilling night; CND1:= chilling night day 1; CND7:= chilling night day 7; CTRL:= control; F6P:= fructose-6-phosphate; FC:= fold change; frc:= fructose; FRCK:= fructokinase; Fv/Fm:= maximum potential quantum efficiency of Photosystem II; G6P:= glucose-6-phosphate; glc:= glucose; GLCK:= glucokinase; nINV:= neutral invertase; md:=midday; NPS:= net photosynthesis; n.s.:= not significant; PC:= permanent cold; suc:= sucrose; SPS:= sucrose phosphate synthase.

**Table S2:**
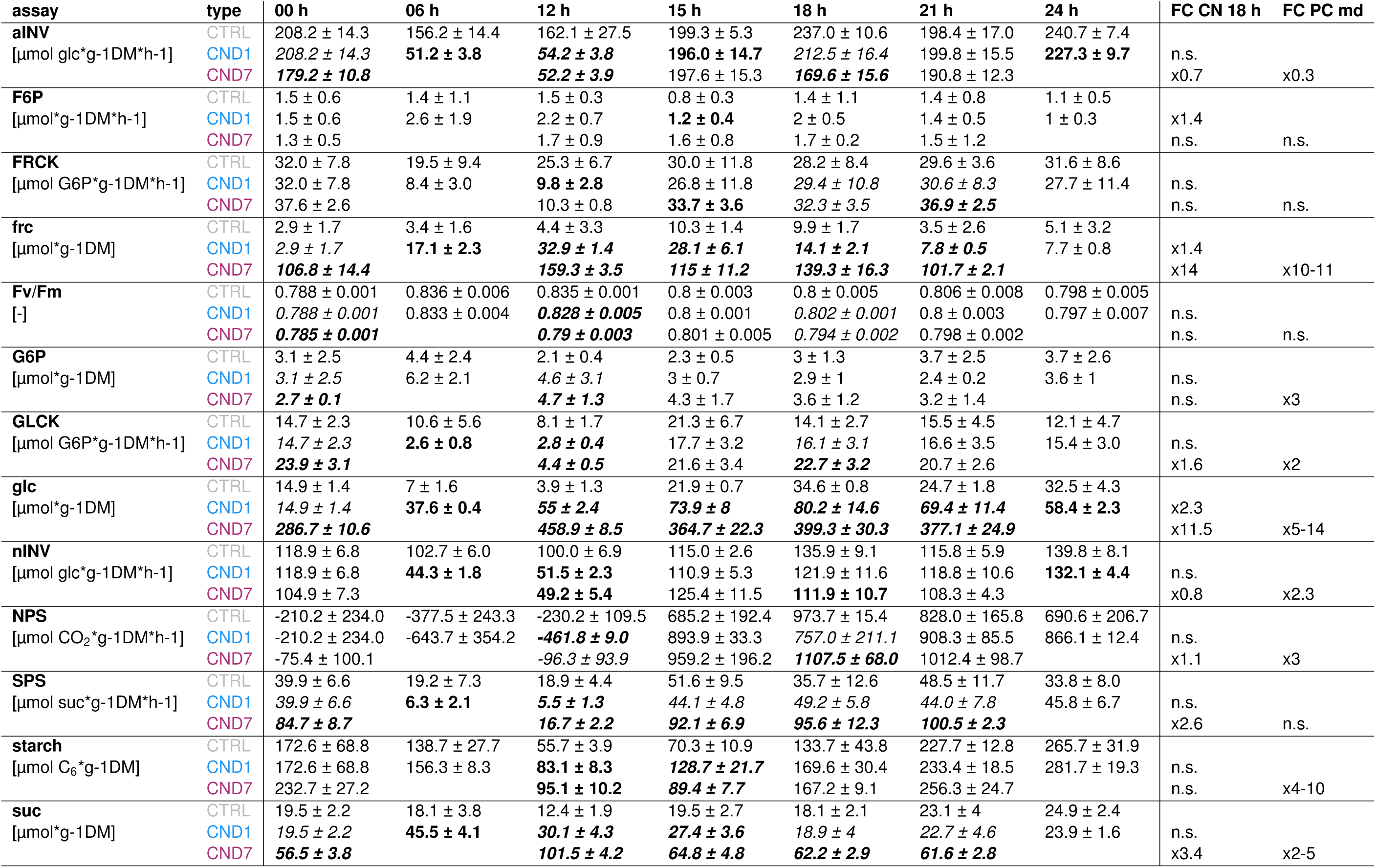
Carbohydrate concentrations and related Arrhenius corrected maximum enzymes activities for control and chilling night conditions. Mean values ± standard deviation for all measured carbohydrate concentrations in [µmol·g^-1^ DM] and related maximum enzyme activities in [µmol·g^-1^ DM·h^-1^]. Maximum enzyme activity of aINV, FRCK, GLCK, nINV and SPS are temperature adapted by Arrhenius correction (Arrhenius, 1889a, 1889b; Laidler, 1984). Significant differences from the control (CTRL) are depicted **bold**. When there is a significant difference between the chilling night day 1 (CND1) and the chilling night day 7 (CND7) they are both depicted in *italics*. FC CN 18 h describes the mean fold change from CTRL to CND7 for the 18 h (midday) time point. FC PC represents mean fold change from midday (md) day 7 of permanent cold condition (4 *^◦^*C) normalized to respective control investigated by Kitashova et al. (2021, 2023). Abbreviations: aINV:= acidic invertase; CN:= chilling night; CND1:= chilling night day 1; CND7:= chilling night day 7; CTRL:= control; F6P:= fructose-6-phosphate; FC:= fold change; frc:= fructose; FRCK:= fructokinase; Fv/Fm:= maximum potential quantum efficiency of Photosystem II; G6P:= glucose-6-phosphate; glc:= glucose; GLCK:= glucokinase; md:=midday; nINV:= neutral invertase; NPS:= net photosynthesis; n.s.:= not significant; PC:= permanent cold; suc:= sucrose; SPS:= sucrose phosphate synthase.

**Table S3:** Fold change carbohydrate concentrations and related maximum enzymes activities (v _max_). Fold change of **A)** not temperature adapted concentrations and maximum enzyme activities. The fold changes were calculated between CND1/7 and the CTRL. **B)** For nINV, FRCK, GLCK, SPS and aINV Arrhenius corrected maximum enzyme activities were used for fold change calculation. Abbreviations: nINV:= neutral invertase; CND1:= chilling night day 1; CND7:= Chilling night day 7; CTRL:= control; F6P:= fructose-6-phosphate; FC:= fold change; frc:= fructose; FRCK:= fructokinase; Fv/Fm:= maximum potential quantum efficiency of Photosystem II; G6P:= glucose-6-phosphate; glc:= glucose; GLCK:= glucokinase; NPS:= Net photosynthesis; suc:= sucrose; SPS:= sucrose phosphate synthase; aINV:= acidic invertase.

| A) |  | B) |  |  |  |  |  |  |  |  |  |  |  |  |  |  |
| --- | --- | --- | --- | --- | --- | --- | --- | --- | --- | --- | --- | --- | --- | --- | --- | --- |
| assay | type | 00 h | 06 h | 12 h | 15 h | 18 h | 21 h | 24 h | type | 0 h | 6 h | 12 h | 15 h | 18 h | 21 h | 24 h |
| nINV<br>[μmol glc*g-1DM*h-1] | CTRL | 1.0 | 1.0 | 1.0 | 1.0 | 1.0 | 1.0 | 1.0 | CTRL | 1.0 | 1.0 | 1.0 | 1.0 | 1.0 | 1.0 | 1.0 |
|  | CND1 | 1.0 | 0.8 | 1.0 | 1.0 | 0.9 | 1.0 | 0.9 | CND1 | 1.0 | 0.4 | 0.5 | 1.0 | 0.9 | 1.0 | 0.9 |
|  | CND7 | 0.9 |  | 1.0 | 1.0 | 1.1 | 0.9 |  | CND7 | 0.9 |  | 0.5 | 1.1 | 0.8 | 0.9 |  |
| F6P<br>[μmol*g-1DM*h-1] | CTRL | 1.0 | 1.0 | 1.0 | 1.0 | 1.0 | 1.0 | 1.0 | CTRL | 1.0 | 1.0 | 1.0 | 1.0 | 1.0 | 1.0 | 1.0 |
|  | CND1 | 1.0 | 1.9 | 1.5 | 1.6 | 1.4 | 1.0 | 0.9 | CND1 | 1.0 | 1.9 | 1.5 | 1.6 | 1.4 | 1.0 | 0.9 |
|  | CND7 | 0.8 |  | 1.2 | 2.0 | 1.2 | 1.1 |  | CND7 | 0.8 |  | 1.2 | 2.0 | 1.2 | 1.1 |  |
| FRCK<br>[μmol F6P*g-1DM*h-1] | CTRL | 1.0 | 1.0 | 1.0 | 1.0 | 1.0 | 1.0 | 1.0 | CTRL | 1.0 | 1.0 | 1.0 | 1.0 | 1.0 | 1.0 | 1.0 |
|  | CND1 | 1.0 | 1.0 | 1.0 | 1.1 | 0.9 | 1.0 | 0.8 | CND1 | 1.0 | 0.4 | 0.4 | 1.1 | 0.9 | 1.0 | 0.9 |
|  | CND7 | 1.2 |  | 1.0 | 1.3 | 1.1 | 1.2 |  | CND7 | 1.2 |  | 0.4 | 1.4 | 1.1 | 1.2 |  |
| frc<br>[μmol*g-1DM] | CTRL | 1.0 | 1.0 | 1.0 | 1.0 | 1.0 | 1.0 | 1.0 | CTRL | 1.0 | 1.0 | 1.0 | 1.0 | 1.0 | 1.0 | 1.0 |
|  | CND1 | 1.0 | 5.0 | 7.5 | 2.7 | 1.4 | 2.2 | 1.5 | CND1 | 1.0 | 5.0 | 7.5 | 2.7 | 1.4 | 2.2 | 1.5 |
|  | CND7 | 36.2 |  | 36.3 | 11.2 | 14.0 | 28.7 |  | CND7 | 36.2 |  | 36.3 | 11.2 | 14.0 | 28.7 |  |
| Fv/Fm<br>[-] | CTRL | 1.0 | 1.0 | 1.0 | 1.0 | 1.0 | 1.0 | 1.0 | CTRL | 1.0 | 1.0 | 1.0 | 1.0 | 1.0 | 1.0 | 1.0 |
|  | CND1 | 1.0 | 1.0 | 1.0 | 1.0 | 1.0 | 1.0 | 1.0 | CND1 | 1.0 | 1.0 | 1.0 | 1.0 | 1.0 | 1.0 | 1.0 |
|  | CND7 | 1.0 |  | 1.0 | 1.0 | 1.0 | 1.0 |  | CND7 | 1.0 |  | 1.0 | 1.0 | 1.0 | 1.0 |  |
| G6P<br>[μmol*g-1DM] | CTRL | 1.0 | 1.0 | 1.0 | 1.0 | 1.0 | 1.0 | 1.0 | CTRL | 1.0 | 1.0 | 1.0 | 1.0 | 1.0 | 1.0 | 1.0 |
|  | CND1 | 1.0 | 1.4 | 2.2 | 1.2 | 1.0 | 0.6 | 1.0 | CND1 | 1.0 | 1.4 | 2.2 | 1.2 | 1.0 | 0.6 | 1.0 |
|  | CND7 | 0.9 |  | 2.2 | 1.9 | 1.2 | 0.9 |  | CND7 | 0.9 |  | 2.2 | 1.9 | 1.2 | 0.9 |  |
| GLCK<br>[μmol G6P*g-1DM*h-1] | CTRL | 1.0 | 1.0 | 1.0 | 1.0 | 1.0 | 1.0 | 1.0 | CTRL | 1.0 | 1.0 | 1.0 | 1.0 | 1.0 | 1.0 | 1.0 |
|  | CND1 | 1.0 | 1.0 | 1.4 | 0.9 | 1.0 | 1.0 | 1.3 | CND1 | 1.0 | 0.3 | 0.3 | 0.9 | 1.0 | 1.1 | 1.2 |
|  | CND7 | 1.6 |  | 2.2 | 1.0 | 1.6 | 1.3 |  | CND7 | 1.6 |  | 0.5 | 1.1 | 1.6 | 1.3 |  |
| glc<br>[μmol*g-1DM] | CTRL | 1.0 | 1.0 | 1.0 | 1.0 | 1.0 | 1.0 | 1.0 | CTRL | 1.0 | 1.0 | 1.0 | 1.0 | 1.0 | 1.0 | 1.0 |
|  | CND1 | 1.0 | 5.3 | 14.2 | 3.4 | 2.3 | 2.8 | 1.8 | CND1 | 1.0 | 5.3 | 14.2 | 3.4 | 2.3 | 2.8 | 1.8 |
|  | CND7 | 19.3 |  | 118.7 | 16.7 | 11.5 | 15.2 |  | CND7 | 19.3 |  | 118.7 | 16.7 | 11.5 | 15.2 |  |
| NPS<br>[μmol CO<sub>2</sub>*g-1DM*h-1] | CTRL | 1.0 | 1.0 | 1.0 | 1.0 | 1.0 | 1.0 | 1.0 | CTRL | 1.0 | 1.0 | 1.0 | 1.0 | 1.0 | 1.0 | 1.0 |
|  | CND1 | 1.0 | 1.7 | 2.0 | 1.3 | 0.8 | 1.1 | 1.3 | CND1 | 1.0 | 1.7 | 2.0 | 1.3 | 0.8 | 1.1 | 1.3 |
|  | CND7 | 0.4 |  | 0.4 | 1.4 | 1.1 | 1.2 | 1.3 | CND7 | 0.4 |  | 0.4 | 1.4 | 1.1 | 1.2 | 1.3 |
| SPS<br>[μmol suc*g-1DM*h-1] | CTRL | 1.0 | 1.0 | 1.0 | 1.0 | 1.0 | 1.0 | 1.0 | CTRL | 1.0 | 1.0 | 1.0 | 1.0 | 1.0 | 1.0 | 1.0 |
|  | CND1 | 1.0 | 1.4 | 1.2 | 0.8 | 1.4 | 0.8 | 1.3 | CND1 | 1.0 | 0.3 | 0.3 | 0.8 | 1.4 | 0.8 | 1.3 |
|  | CND7 | 2.2 |  | 3.8 | 1.8 | 2.7 | 2.1 |  | CND7 | 2.2 |  | 1.0 | 1.8 | 2.7 | 2.1 |  |
| starch<br>[μmol C<sub>6</sub>*g-1DM] | CTRL | 1.0 | 1.0 | 1.0 | 1.0 | 1.0 | 1.0 | 1.0 | CTRL | 1.0 | 1.0 | 1.0 | 1.0 | 1.0 | 1.0 | 1.0 |
|  | CND1 | 1.0 | 1.1 | 1.5 | 1.8 | 1.3 | 1.0 | 1.0 | CND1 | 1.0 | 1.1 | 1.5 | 1.8 | 1.3 | 1.0 | 1.0 |
|  | CND7 | 1.3 |  | 1.7 | 1.3 | 1.3 | 1.1 |  | CND7 | 1.3 |  | 1.7 | 1.3 | 1.3 | 1.1 |  |
| suc<br>[μmol*g-1DM] | CTRL | 1.0 | 1.0 | 1.0 | 1.0 | 1.0 | 1.0 | 1.0 | CTRL | 1.0 | 1.0 | 1.0 | 1.0 | 1.0 | 1.0 | 1.0 |
|  | CND1 | 1.0 | 2.5 | 2.4 | 1.4 | 1.0 | 1.0 | 1.0 | CND1 | 1.0 | 2.5 | 2.4 | 1.4 | 1.0 | 1.0 | 1.0 |
|  | CND7 | 2.9 |  | 8.2 | 3.3 | 3.4 | 2.7 |  | CND7 | 2.9 |  | 8.2 | 3.3 | 3.4 | 2.7 |  |
| aINV<br>[μmol glc*g-1DM*h-1] | CTRL | 1.0 | 1.0 | 1.0 | 1.0 | 1.0 | 1.0 | 1.0 | CTRL | 1.0 | 1.0 | 1.0 | 1.0 | 1.0 | 1.0 | 1.0 |
|  | CND1 | 1.0 | 0.9 | 1.0 | 1.0 | 0.9 | 1.0 | 1.0 | CND1 | 1.0 | 0.3 | 0.3 | 1.0 | 0.9 | 1.0 | 0.9 |
|  | CND7 | 0.8 |  | 0.9 | 1.0 | 0.7 | 1.0 |  | CND7 | 0.9 |  | 0.3 | 1.0 | 0.7 | 1.0 |  |

**Table S4:**
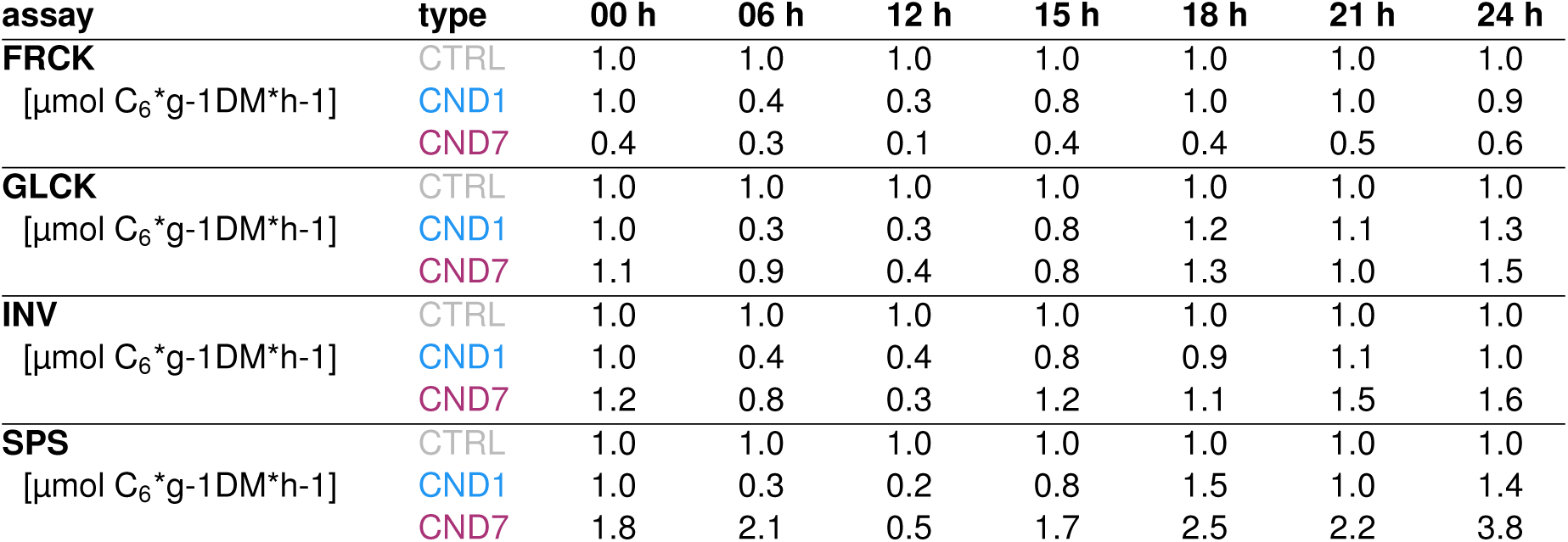
Fold change of modeled enzymes activities *v* _pred_. The mean fold changes were calculated between CND1 or CND7 normalized to the CTRL. Abbreviations: CND1:= chilling night day 1; CND7:= Chilling night day 7; CTRL:= control; FRCK:= fructokinase; GLCK:= glucokinase; INV:= invertase; SPS:= sucrose phosphate synthase;

#### A.3.9 Growth under Chilling Night Conditions

**Figure S16:**
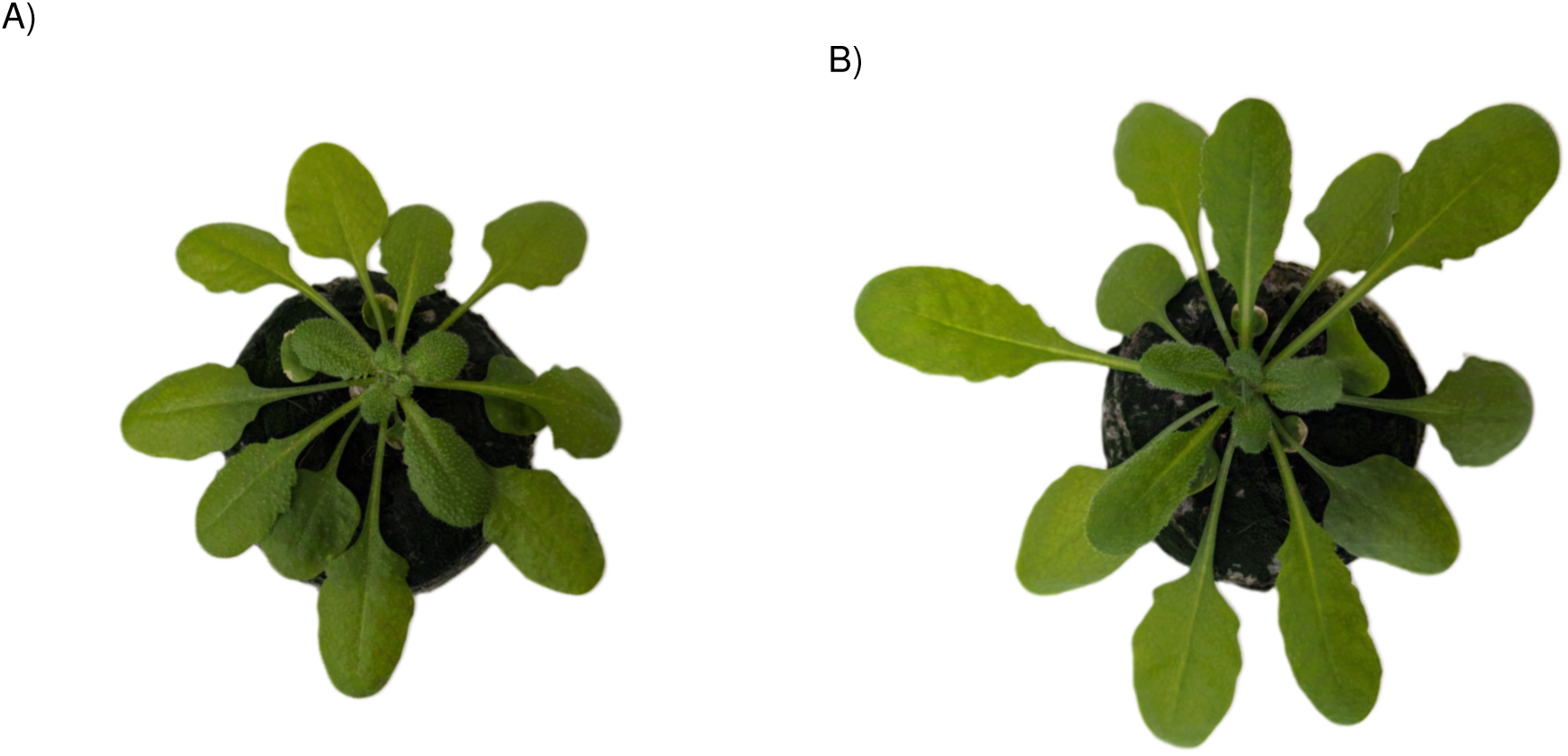
Leaf growth is not reduced by chilling nights in Col-0. **A)** Col-0 after first chilling night (no visible difference to control). **B)** Col-0 after seven chilling nights. Scale: jiffy pellet (7): 44 mm.

#### A.3.10 Modeled Enzyme Rates

**Figure S17:**
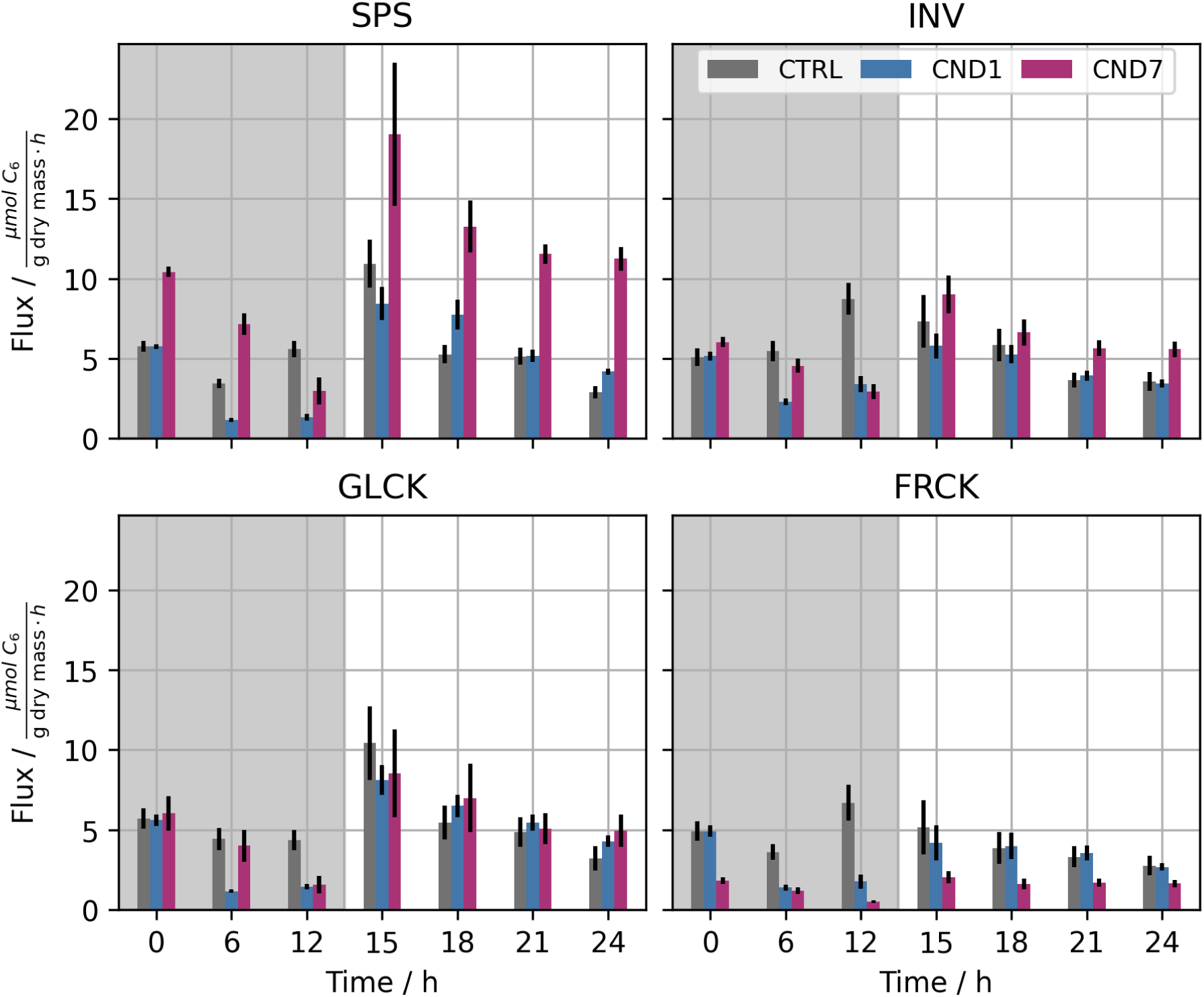
Fluxes of key enzymes across experimental conditions. Predicted *v* _pred_ as estimated by an ensemble of 100 augmented neural ODE (ANODE) models in i) control (CTRL, gray, 18 *^◦^*C night and 22 *^◦^*C day), ii) chilling night day one (CND1, blue, 4 *^◦^*C night and 22 *^◦^*C day) and iii) chilling night day seven (CND7, magenta, 4 *^◦^*C nights and 22 *^◦^*C days). *v* _pred_ in µmol C_6_ equivalents per gram DM per hour are plotted against the TPs [h]. Predictions below the 0.2 and above the 0.8 quantile per condition and time point removed for visual clarity. Error bars indicate mean *±* standard deviation across the ensemble. Abbreviations: CND1:= chilling night day 1; CND7:= chilling night day 7; CTRL:= control; FRCK:= fructokinase; GLCK:= glucokinase; INV:= invertase; SPS:= sucrose phosphate synthase.

#### A.3.11 Cumulative Enzyme Rates per Night and Day

**Figure S18:**
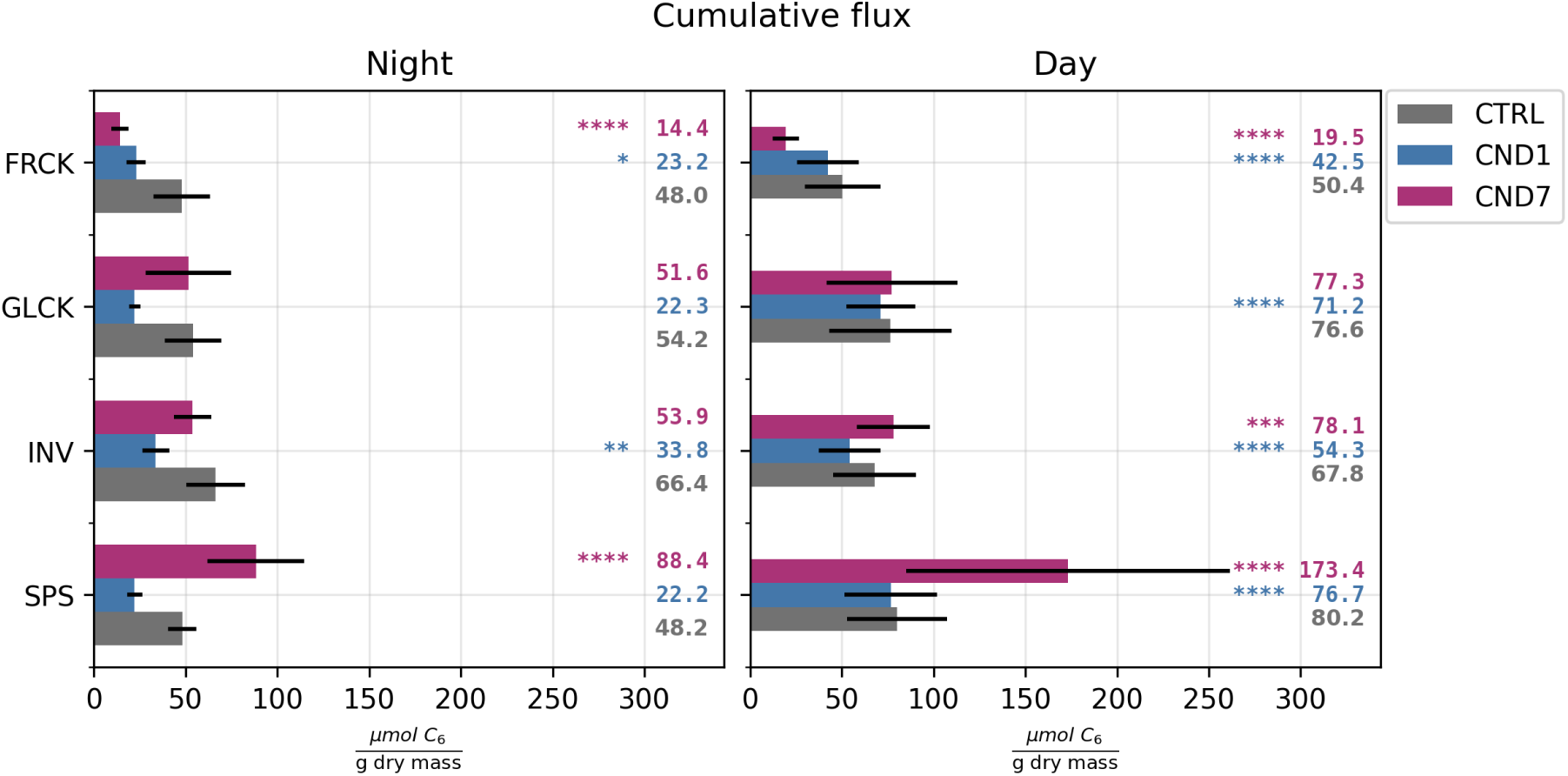
Cumulative predicted ensemble fluxes partitioned into daytime and nighttime contributions. Bars represent ensemble mean values in i) control (CTRL, gray, 18 *^◦^*C night and 22 *^◦^*C day), ii) chilling night day one (CND1, blue, 4 *^◦^*C night and 22 *^◦^*C day) and iii) chilling night day seven (CND7, magenta, 4 *^◦^*C nights and 22 *^◦^*C days), with error bars indicating *±* standard deviation. Cumulative fluxes were computed via trapezoidal integration over the respective time steps and predictions below the 0.2 and above the 0.8 quantile per condition and time point removed for visual clarity, but retained for statistical analysis. Statistical significance between CTRL and each treatment condition (CND1, CND7) was assessed using two-sided Mann–Whitney U tests, with p-values corrected for multiple comparisons using the Benjamini–Hochberg false discovery rate method. Significance levels are denoted as: *∗p <* 0.05, *∗ ∗ p <* 0.01, *∗ ∗ ∗p <* 0.001, *∗ ∗ ∗ ∗ p <* 0.0001. Abbreviations: CND1:= chilling night day 1; CND7:= chilling night day 7; CTRL:= control; FRCK:= fructokinase; GLCK:= glucokinase; INV:= invertase; SPS:= sucrose phosphate synthase.

#### A.3.12 Metabolite Fits

**Figure S19:**
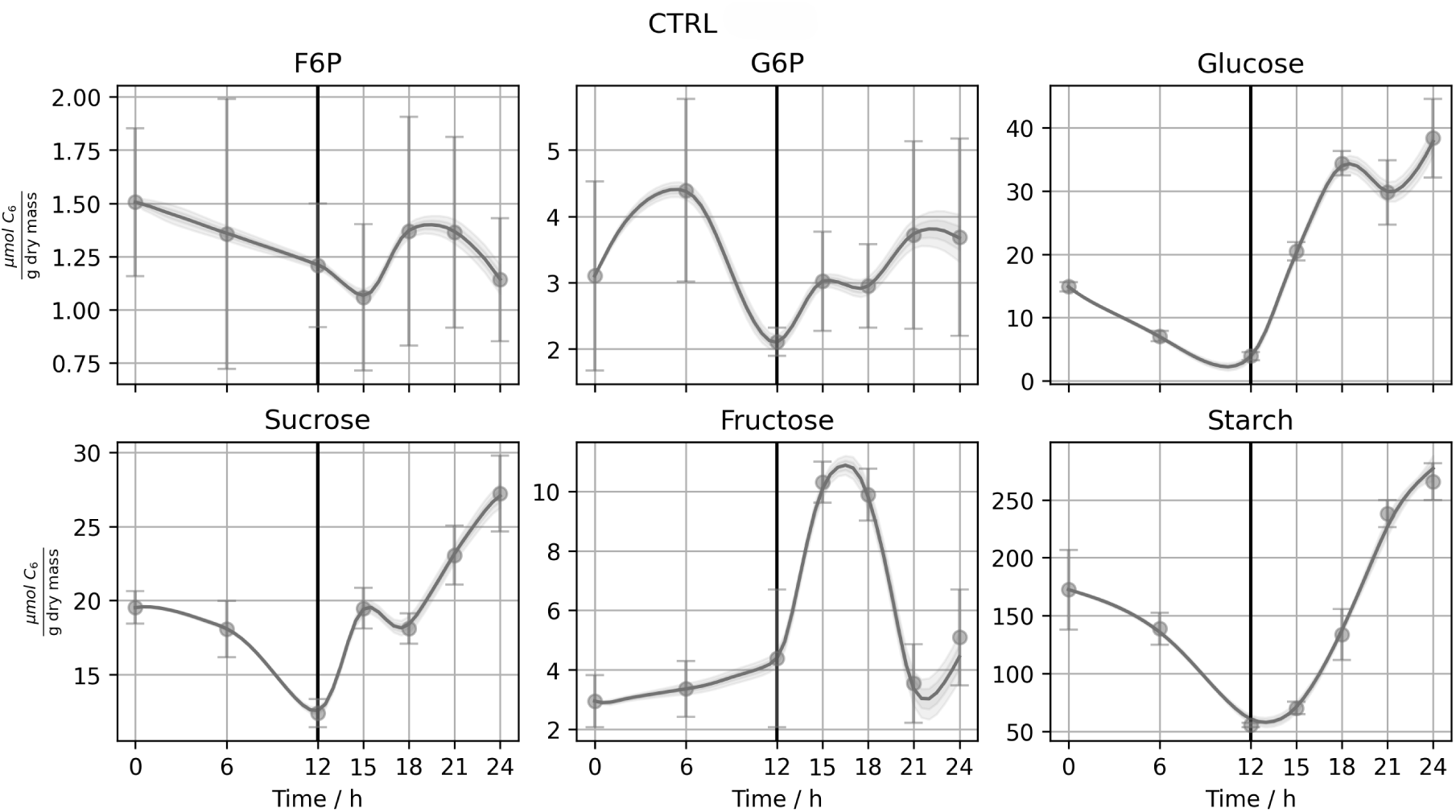
Time series predictions of key intermediate concentrations generated by an ensemble of augmented neural ODE (ANODE) models for control condition (CTRL). Each panel shows a single intermediate species. Solid lines represent the ensemble mean prediction, and shaded regions indicate *±* standard deviation across the ensemble with predictions below the 0.2 and above the 0.8 quantile per condition and time point removed for visual clarity. Measured concentrations are overlaid as scatter points. Abbreviations: CTRL:= control; F6P:= fructose-6-phosphate; G6P:= glucose-6-phosphate.

**Figure S20:**
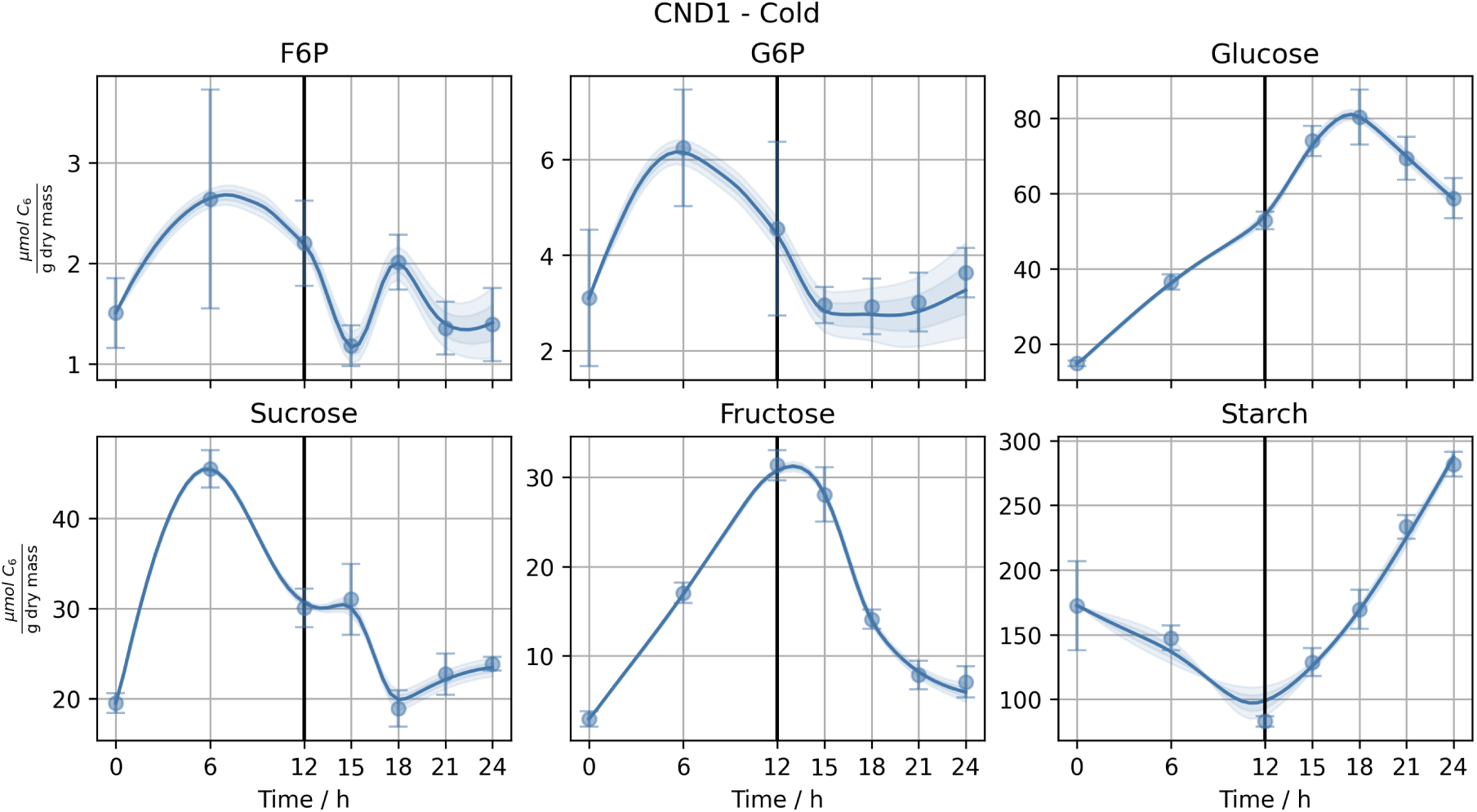
Time series predictions of key intermediate concentrations generated by an ensemble of augmented neural ODE (ANODE) models for cold night condition on day 1 (CND1). Each panel shows a single intermediate species. Solid lines represent the ensemble mean prediction, and shaded regions indicate *±* standard deviation across the ensemble with predictions below the 0.2 and above the 0.8 quantile per condition and time point removed for visual clarity. Measured concentrations are overlaid as scatter points. Abbreviations: CND1:= chilling night day 1; F6P:= fructose-6-phosphate; G6P:= glucose-6-phosphate.

**Figure S21:**
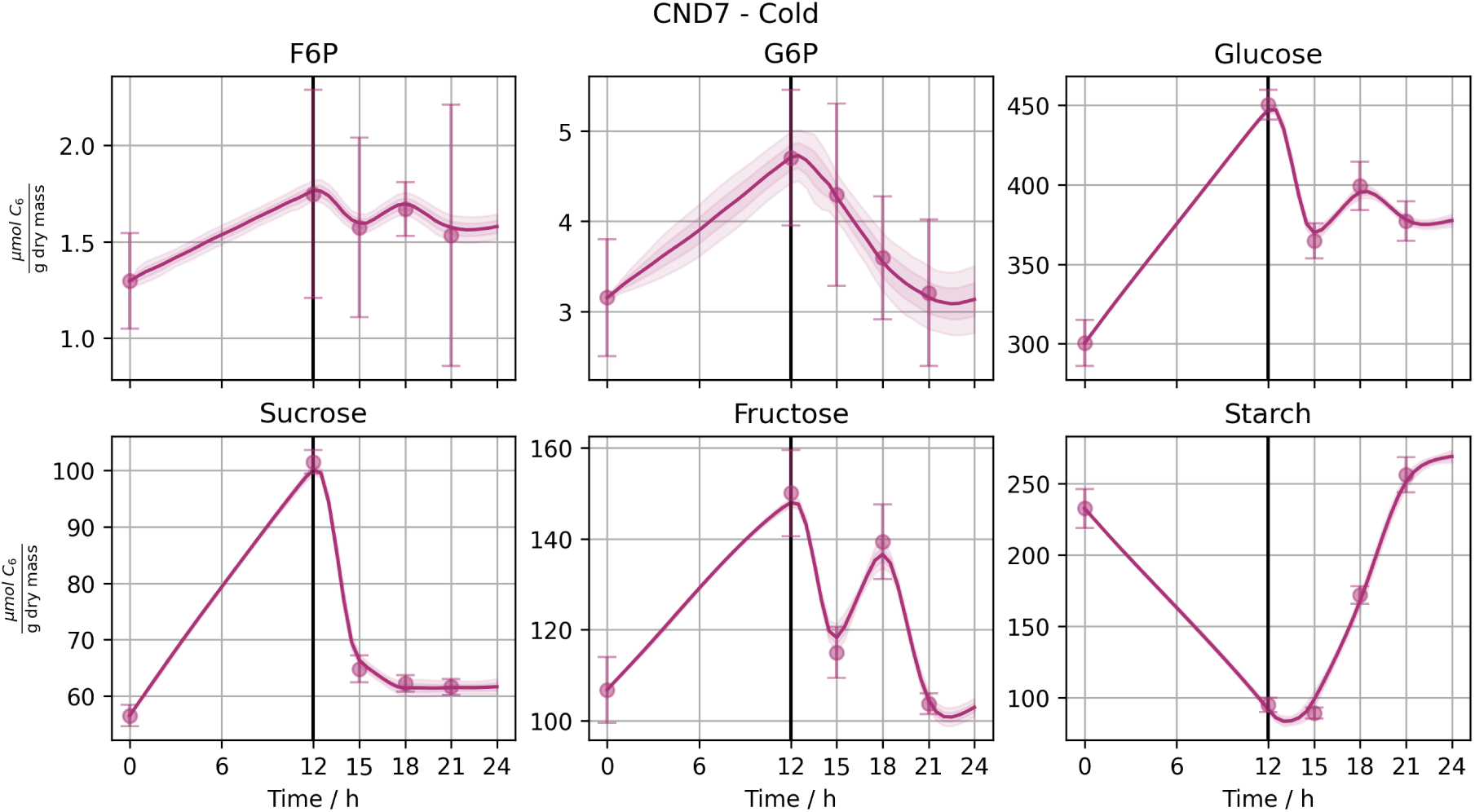
Time series predictions of key intermediate concentrations generated by an ensemble of augmented neural ODE (ANODE) models for cold night condition on day 7 (CND7). Each panel shows a single intermediate species. Solid lines represent the ensemble mean prediction, and shaded regions indicate *±* standard deviation across the ensemble with predictions below the 0.2 and above the 0.8 quantile per condition and time point removed for visual clarity. Measured concentrations are overlaid as scatter points. Abbreviations: CND7:= chilling night day 7; F6P:= fructose-6-phosphate; G6P:= glucose-6-phosphate.

## Notes

### Competing Interest Statement

The authors have declared no competing interest.

